# Single cell analysis of dup15q syndrome reveals developmental and postnatal molecular changes in autism

**DOI:** 10.1101/2023.09.22.559056

**Authors:** Yonatan Perez, Dmitry Velmeshev, Li Wang, Matthew White, Clara Siebert, Jennifer Baltazar, Natalia Garcia Dutton, Shaohui Wang, Maximilian Haeussler, Stormy Chamberlain, Arnold Kriegstein

## Abstract

Duplication 15q (dup15q) syndrome is the most common genetic cause of autism spectrum disorder (ASD). Due to a higher genetic and phenotypic homogeneity compared to idiopathic autism, dup15q syndrome provides a well-defined setting to investigate ASD mechanisms. Previous bulk gene expression studies identified shared molecular changes in ASD. However, how cell type specific changes compare across different autism subtypes and how they change during development is largely unknown. In this study, we used single cell and single nucleus mRNA sequencing of dup15q cortical organoids from patient iPSCs, as well as post-mortem patient brain samples. We find cell-type specific dysregulated programs that underlie dup15q pathogenesis, which we validate by spatial resolved transcriptomics using brain tissue samples. We find degraded identity and vulnerability of deep-layer neurons in fetal stage organoids and highlight increased molecular burden of postmortem upper-layer neurons implicated in synaptic signaling, a finding shared between idiopathic ASD and dup15q syndrome. Gene co-expression network analysis of organoid and postmortem excitatory neurons uncovers modules enriched with autism risk genes. Organoid developmental modules were involved in transcription regulation via chromatin remodeling, while postmortem modules were associated with synaptic transmission and plasticity. The findings reveal a shifting landscape of ASD cellular vulnerability during brain development.

---

Autism spectrum disorder (ASD) is a complex neurodevelopmental and neurological disorder with a strong genetic contribution ^1^. Despite a heterogeneity of clinical manifestations and underlying genetics, systems biological and functional genomic studies of the brains of ASD patients suggest convergence of disease pathology on common pathways. These include alternative splicing, synaptic signaling, microglia activation, epigenetic modifications, and neuron-specific transcriptional regulation ^1–10^. Although many studies indicate that some ASD-associated gene modules are co-expressed during prenatal development (early to mid-fetal), other studies highlight ASD-associated genes expressed during postnatal development ^5–11^. Recently, organoid models of the developing human cerebral cortex have enabled identification of the cell-type-specific pathology of monogenic ASD syndromes ^12–14^, and may prove pivotal in the future to further our knowledge of early developmental ASD-associated phenotypes. Postnatally, specific brain regions underlying higher-order cognitive processes, such as the prefrontal and temporal cortex and the limbic system, have been demonstrated to be specifically affected in ASD ^6,8,15–17^. In addition to idiopathic cases, ASD is also associated with several genetic syndromes. The most common is 15q11-q13 duplication syndrome (dup15q syndrome), accounting for up to 3% of all ASD cases ^18^. Dup15q is commonly caused by a *de-novo* maternally derived duplication of the Prader-Willi/Angelman critical region (PWACR), typically encompassing tens of genes (depending on the DNA breaking point). This genomic locus contains both maternally and paternally imprinted genes, resulting in monoallelic expression that is necessary for normal neurodevelopment. This type of expression is controlled by the differentially methylated bipartite Prader-Willi syndrome imprinting center (PWS-IC) and the Angelman syndrome imprinting center (AS-IC) ^19^. Loss of paternally expressed genes results in Prader-Willi syndrome (PWS, OMIM #176270), while maternal gene deficiency causes Angelman syndrome (AS, OMIM #105830); both are phenotypically distinct from dup15q syndrome. The duplication mainly occurs in two forms, either as an extra isodicentric 15 chromosome idic(15q), or as an interstitial duplication int.dup(15q). Most dup15q patients meet the criteria for ASD. Other patient characteristic phenotypes include hypotonia, variable degrees of intellectual disability (ID), motor delay, and epilepsy. In this study, we aim to gain insight into the cell-type specific developmental and postnatal molecular changes of dup15q syndrome and explore potential dysregulated programs that converge between dup15q and idiopathic ASD. We use single-nucleus RNA sequencing (snRNAseq) of dup15q postmortem samples from three different cortical regions, as well as single cell RNA sequencing (scRNAseq) of iPSC-derived cortical organoids, to model dup15q associated transcriptional changes throughout development and in adulthood.

## Comprehensive single-cell molecular profiling of dup15q

We performed nuclei isolation and snRNA-seq of 49 snap frozen post-mortem tissue samples from the prefrontal (PFC), temporal (TC) and anterior cingulate (ACC) cortical regions of 11 dup15q patients and 17 neurotypical controls (**Fig. 1a**). All dup15q patients were diagnosed with ASD. Subjects ranged from 8 to 39 years old, and both dup15q and control samples were matched for age, sex, RNA integrity number (RIN), and post-mortem interval (**Extended Data Fig. 1a-e; Extended Data Table 1**). Sections of snap-frozen cortical blocks were collected from the entire span of cortical grey matter and adjacent subcortical white matter. Tissue was lysed and nuclei were extracted using ultracentrifugation in a sucrose gradient. Nuclei capture and library preparation were done using the 10x Genomics platform. In total, we generated 345,861 single-nuclei gene expression profiles that passed our quality control standards (methods); 145,346 from dup15q subjects and 200,515 from neurotypical controls (**Fig. 1a**).

**Figure 1.**
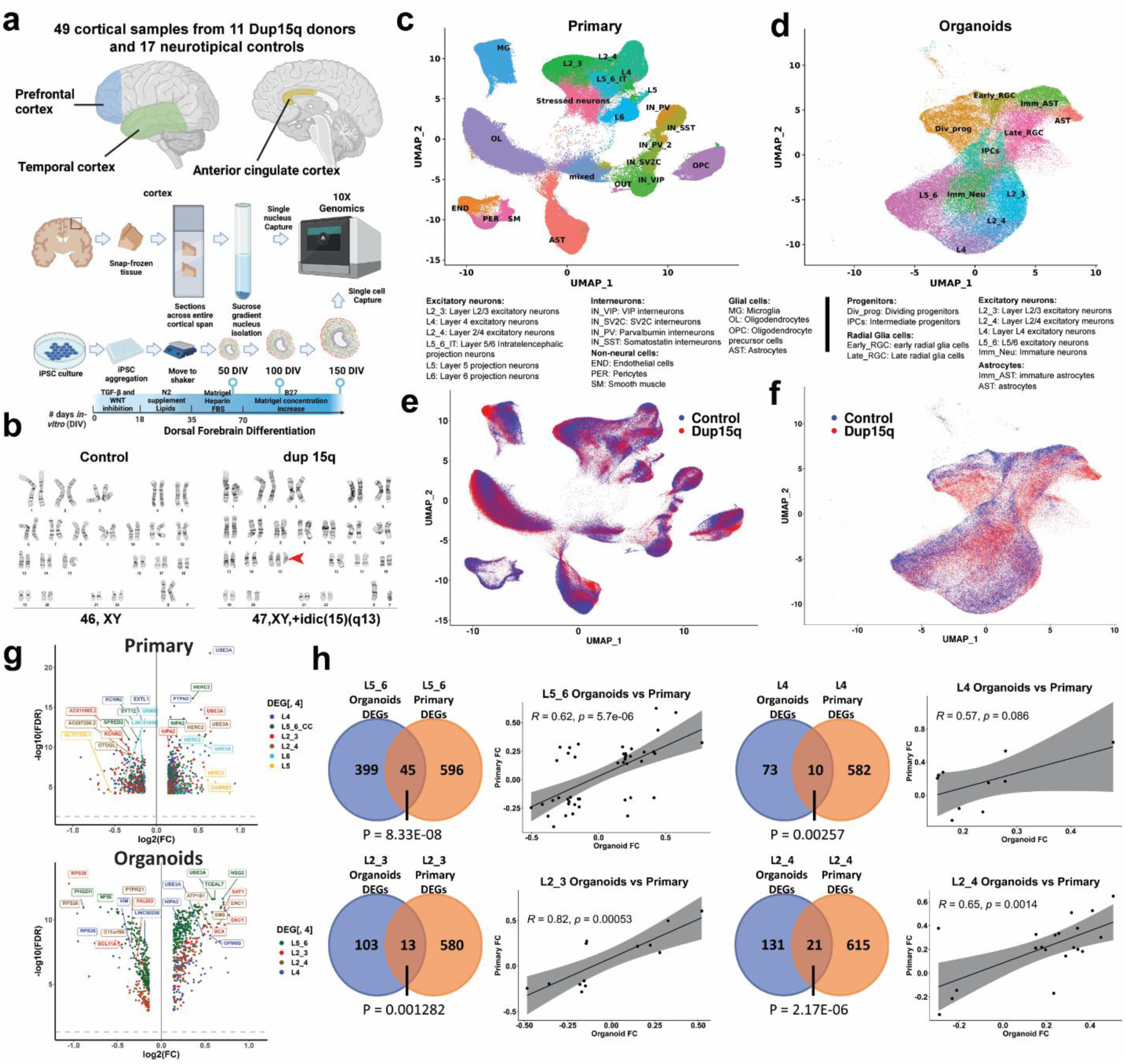
Comprehensive single-cell molecular profiling of dup15q syndrome using postmortem cortical samples and cortical organoids. **a)** Illustration of experimental design, sample collection and cell capture. **b)** G-banding karyotype of normal and idic(15q) iPSC lines. **c)** Unbiased clustering of single nuclei and annotated cell types of postmortem samples. **d)** Unbiased clustering of cortical organoid single cells and annotated cell types. **e)** Primary nuclei clustered by genotype, showing equal contribution of dup15q and control samples to all cell types. **f)** Organoid cells clustered by genotype, showing similar contributions from dup15q and control organoids to all cell types. **g)** Volcano plots for cell type– specific genes differentially expressed in both primary and organoid excitatory neurons. **h)** Overlap between DEGs of primary and organoid excitatory neurons as well as Pearson’s correlation coefficient of shared genes fold changes.

For cortical organoids, we used three patient-derived dup15q (two idic(15q) and one interstitial triplication) and three control human induced pluripotent (hiPSC) stem cell lines (**Fig. 1b, Extended Data Table 1**). All lines underwent karyotype analysis prior to cortical organoid differentiation (**Extended Data Fig. 1f**). Cortical organoids were generated using a dorsal forebrain differentiation protocol ^20,21^, and cultures from different lines demonstrated expected and comparable differentiation progression throughout sampling timepoints and across genotypes (**Extended Data Fig. 1g**). Single cells were captured at 50, 100 and 150 days of *in vitro* differentiation (DIV) using the 10x genomics platform, roughly corresponding to, peak neurogenic and gliogenic stages of early second trimester *in utero* development ^20,22^. Using this approach, we were able to generate 106,302 expression profiles from organoid cells: 63,500 from dup15q and 42,802 from controls, across three *in vitro* developmental timepoints (**Fig. 1a**).

We detected a median of 3108 genes and 7103 transcripts per primary nuclei, and a median of 2672 genes and 7908 transcripts per organoid cell. In both datasets we observed glutamatergic neurons expressing higher numbers of genes and transcripts compared to other cell types (**Extended Data Fig. 2a-b**). Dimensionality reduction and unbiased clustering of postmortem nuclei from all cortical regions and organoid cell profiles were done separately (**Fig. 1c-d**). Clusters were annotated based on the expression of cell type markers (**Extended Data Fig. 2c-d**) for molecularly defined cell types annotated in the Allen Brain Atlas ^23^. After removing a cluster of neuronal debris (**Methods, Extended Data Fig. S2e-g**), we identified 17 specific cell types for primary nuclei (ten neuronal, four glial and three non-neural cell types) and 11 for cortical organoids. Primary nuclei clusters included known subtypes of excitatory neurons, including subcortical and corticocortical projection neurons, known subtypes of both MGE and CGE-derived inhibitory neurons, as well as vascular and glial cell subtypes (**Fig. 1c**). Organoid clustering identified two radial glia populations (early RGCs and late RGCs). Early RGCs were dominant in 50 DIV organoids and expressed *SOX2*, *HES1*, *EMX1, PALLD and PDGFD*, consistent with ventricular radial glia identity ^24^. Late RGCs cells were mostly present in organoids at 100-150 DIV and expressed higher levels of *HOPX*, *PEA15* and *PTPRZ1*, consistent with outer radial glia identity ^24^ (**Extended Data Fig. 2g**). We also found dividing and intermediate progenitors as well as known subtypes of excitatory neurons with identity matching specific cortical layers (**Fig. 1d, Extended Data Fig. 2c**). In addition, we found two clusters of GABAergic interneurons expressing *GAD2*, *DLX2*, *DLX5, DLX-AS1* and lacking *NEUROD2/6* expression, as previously reported ^12,13,25,26^. These were subdivided into early GABAergic neurons (mostly arising from organoids at 50 DIV) with a lateral ganglionic eminence (LGE)-like identity, expressing *SIX3*, *MEIS2* and *PBX3,* and a late interneuron identity (more prominent in organoids at 100 and 150 DIV) with a putative caudal ganglionic eminence (CGE)-like origin expressing *SCGN*, *SP8*, *CALB2* and *PROX1* (**Extended Data Fig. 3a**) ^27,28^. However, these clusters were present in only a subset of the lines (one control and two mutant) and were thus removed from further analysis (**Extended Data Fig. 3b**). Nuclei captured from different samples and from all cortical regions were well intermixed, contributing to all cell types. The cell contribution from different organoids was uniform as well, indicating a lack of strong batch effect (**Figs. 1c,1e, Extended DataFig. 3c**). Organoid cells clustered by days of *in vitro* differentiation demonstrated the sequential emergence and differentiation of excitatory neurons and astrocytes, comparable to human *in-utero* cortex development ^28^ (**Extended Data Fig. 1g, 3d**). For example, organoids sampled at 50 DIV mostly contained early radial glia, intermediate progenitors, and deep-layer (DL) projection neurons. At 100 DIV, organoids mostly contained immature neurons, DL neurons, and immature astrocytes. Organoids captured at 150 DIV contained late radial glia cells, upper-layer (UL) neurons, and mature astrocytes (**Extended Data Fig. 3e**). The fraction of dup15q and control nuclei contributing to each cell type was relatively uniform across cell types. As expected, cell contribution was more heterogeneous and variable between organoids from different lines (**Extended Data Fig. 3f**), likely due to the influence of individual line-specific genomic contexts ^13^.

## Vulnerability of dup15q DL excitatory neurons throughout development

To identify cell-type specific differential gene expression changes between dup15q and controls in both organoids and primary tissue, we utilized a linear mixed model (LMM) approach that controls for confounders (Methods). In primary tissue, we identified 7,896 differential expression events (q value < 0.05; expression fold change ≥ 14%) in 3,634 unique genes across all cell types. Of these, 4,809 genes were upregulated (∼61%) and 3,087 were downregulated (∼39%) **(Extended Data Table 2)**. In Organoids, we identified 2,914 differentially expressed genes (DEGs) in 1,657 unique genes across cell types. Of these, 1,341 were upregulated (∼46%) and 1,573 were downregulated (∼54%) **(Extended Data Table 3)**. We used RNA *in situ* hybridization to validate some of the most differentially expressed genes across all cell types in brain tissue sections **(Extended Data Figs. 4a-c)**. Genes within the 15q11.2-q13.1 duplicated locus (such as *UBE3A*, *GABRB5*, *GABARG3*, *NIPA2* and *HERC2*), were among the highest and most recurrently overexpressed genes across all cell types in both organoids and primary tissue (**Fig. 1g, Extended Data Tables 2,3**). Consistent with previous observations ^18^, when we intersected all primary and organoid overexpressed genes with the list of genes within the putative duplicated locus (15q11.2-q13.1), we found 21 duplicated genes overlapping with primary upregulated genes (**P = 1.15-E06, hypergeometric test; Extended Data Fig. 5a**) and 13 with organoid upregulated genes (**P=1.58-E07, hypergeometric test; Extended Data Fig.5b**). That most of the differentially expressed genes are located outside of the duplicated region indicates that the majority of dup15q gene dysregulation is not driven by DNA dose-dependent expression, but rather via indirect mechanisms regulating gene expression. Several duplicated genes were most differentially expressed specifically in excitatory neurons (**Fig. 1g**), suggesting cell type-specific mechanisms that can modulate expression of genes in the 15q11.2-q13.1 duplicated locus. Despite developmental stage differences, technical differences, and overall differences in gene expression between primary and organoid cell types (**Extended Data Fig. 5c**), we found that all excitatory neurons had statistically significant DEG overlap between organoids and primary tissue. (P < 0.05, hypergeometric test; **Fig. 1h**). This is not surprising considering the overexpression of duplicated genes. Nonetheless, we found that DL neurons had the most overlapping DEGs between organoids and primary cells (45 genes, accounting for >10% of all organoid DEGs and 7% of primary corresponding cells; P = 8.3E-08, hypergeometric test) while having the lowest proportion of overlapping duplicated genes (only ∼11% of overlapping DEGs were duplicated genes). All excitatory neurons, aside from L4 neurons (P = 0.086), showed a significant statistical correlation between expression fold changes of overlapping dup15q associated genes (Pearson’s P value < 0.05; **Fig. 1h, Extended Data Table 3**).

To test the degree by which dup15q associated transcriptional changes affect each of the cell types, we performed gene burden analysis (Methods). In primary tissue, we found that all excitatory neurons were highly affected. Specifically, UL projection neurons (L2_3) had the largest number of DEGs followed by DL intratelencephalic (IT) projecting neurons (L5_6_IT). In organoids, early radial glia had the largest number of DEGs, followed by DL (L5_6) projection neurons (**Fig. 2a**). This is in line with previous studies demonstrating a key point of convergence for ASD implicating both DL and UL glutamatergic neurons ^5,7,8^. The burden discrepancy between organoid cells and primary nuclei might be attributed to developmental changes captured by our experimental paradigm. It is likely that dup15q associated molecular changes of UL neurons occur later in development and are not yet manifest in the relatively immature UL neurons in our organoids ^22^. In contrast, DL neurons arise early during organoid development and are thus relatively more mature and more susceptible to molecular changes at this stage. These findings provide evidence for the vulnerability of glutamatergic cortical neurons in dup15q syndrome across developmental stages.

**Figure 2.**
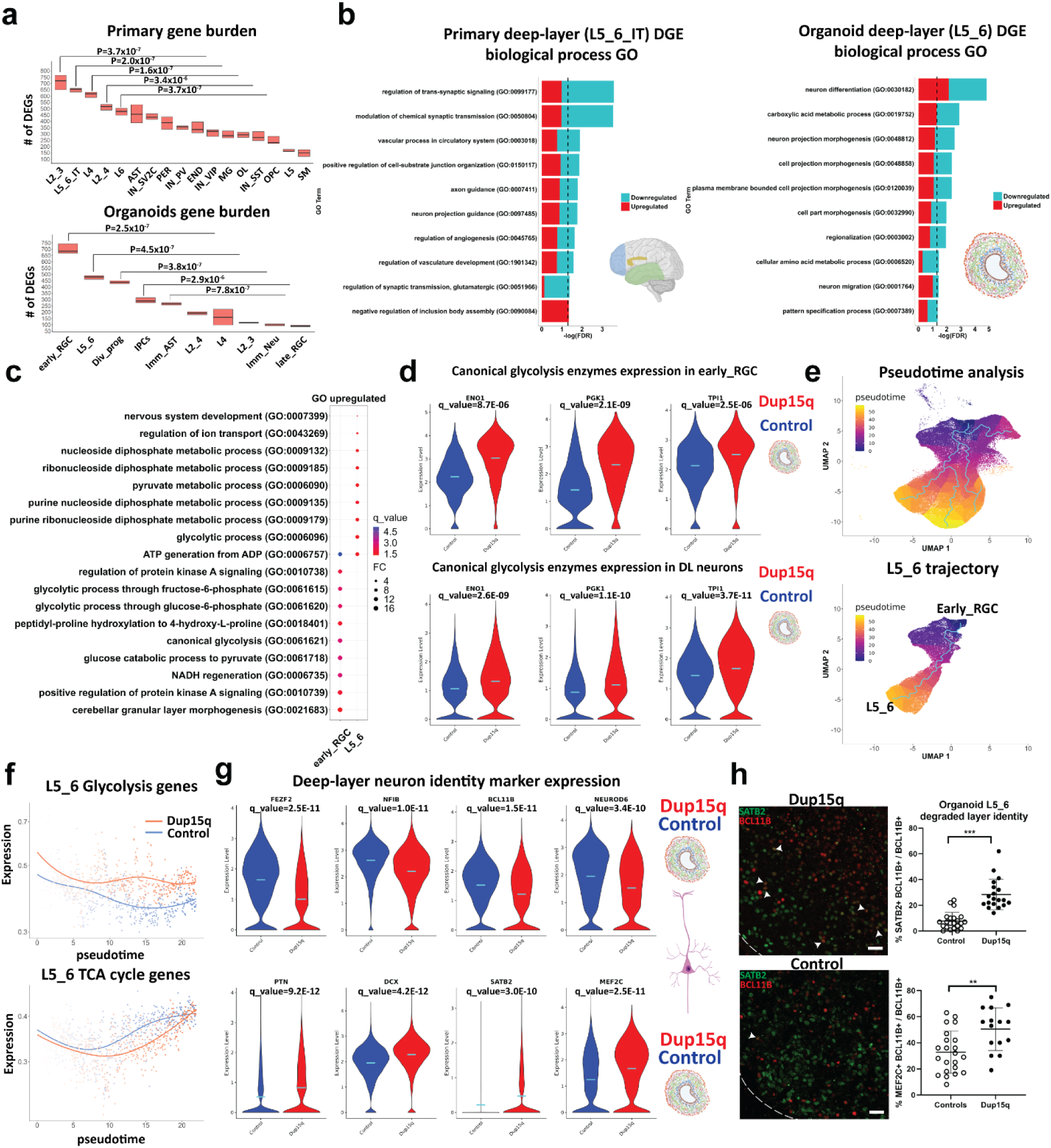
increased glycolysis in dup15q cortical organoids lead to degraded deep-layer neuronal identity. **a)** Gene burden analysis of primary and cortical organoid cells-types. (Mann-Whitney U test). **b)** GO analysis of primary and cortical deep-layer (DL) excitatory neurons. **c)** GO analysis of organoid early RGC and DL neurons showing enrichment of glycolysis associated terms (q value = -log(FDR); FC = Fold change). **d)** Violin plots showing increased expression of canonical glycolytic enzymes in dup15q early RGCs and DL neurons. **e)** Pseudotime analysis identifies organoid trajectories comparable to *in vivo* cortical development (top panel). Identifying dynamic gene expression changes along the DL neuron trajectory (bottom panel). **f)** DL neurons of dup15q syndrome cortical organoids are expressing high levels of glycolysis genes and attenuated TCA cycle genes along pseudotime. **g)** Violin plots showing co-expression of DL markers, immature neuron markers, and UL neuron markers, indicating degraded identity of dup15q organoid DL neurons. (Control in blue and dup15q in red). **f)** IF and Co-expression analysis of organoid DL and UL neuron markers (scale bar = 100µm; white arrowheads indicating BLC11B and SATB2 co-expression).

Among some of the prominent DEGs shared between organoids and primary DL cortical neurons are the duplicated genes *UBE3A* and *GABRB3*, thought to be among the major drivers of autism and epilepsy phenotypes respectively. Additional DL overlapping genes are known ASD-risk genes such as the BAF (SWI/SNF) chromatin remodeling complex associated transcription factor, *BCL11A,* and the netrin receptor, *UNC5A*, required for axon guidance. Gene ontology (GO) analysis of differentially expressed genes, identified regulation of neuron projection as a common associated biological process that is affected in both organoids and primary dup15q DL neurons (**Fig. 2b**). These findings highlight temporal differences of cell type vulnerability and indicate dysregulated cell-type specific gene expression in dup15q. Together, this uncovers vulnerability of DL projection neurons, implicated in neuron projection, which is not entirely driven by duplicated genes, and highlights a common cell-type specific biological defect in dup15q that persists throughout development.

## Increased glycolysis in dup15q progenitors leads to degraded neuronal layer identity

Organoid dup15q early radial glia and DL neurons share many DEGs (P = 8.485E-45, hypergeometric test, **Extended Data Table 3**). Gene ontology analysis of overexpressed genes identified glycolysis as a common molecular pathway in both cell types, reflecting overexpression of canonical glycolytic enzymes such as *ENO1*, *TPI1* and *PGK1* (**Figs. 2c-d, Extended Data Fig. 5d**). Glycolysis and mitochondrial dynamics are tightly regulated and are critical for proper forebrain development. The transition from hypoxic, mostly anerobic glycolysis-dependent, proliferative states to more bioenergetically demanding oxidative phosphorylation (OXPHOS) reliant differentiation states has been well studied ^29^. In addition, an increase in oxygen consumption, reliance on the tricarboxylic acid (TCA) cycle and shutoff of anerobic glycolysis was shown to be critical for differentiation and maturation of human cultured neurons ^30^. Thus, cellular metabolism program shifts may be important for cell fate transition and cellular identity ^31,32^ during cortical development. Furthermore, cellular metabolic states and their dysregulation may cause Intellectual disability, epilepsy, and neuropsychiatric disorders including autism ^33–36^. To test whether glycolysis enrichment is unique to dup15q cells transitioning from radial glia to DL neurons, we performed pseudotime analysis using Monocle 3 in control and dup15q organoids (Methods). We inferred organoid trajectories that faithfully recapitulated *in vivo* lineage progression (**Fig. 2e**) and identified genes that are differentially expressed between control and dup15q organoids in cell-specific lineages along pseudotime **(Extended Data Table 4)**. We then plotted the expression of glycolysis, TCA cycle, and OXPHOS genes (hallmark and KEGG genes, MSigDB dataset; Methods) along pseudotime for each unique cell-specific trajectory. We found enrichment in glycolysis concomitant with attenuated TCA cycle expression only within the dup15q DL neuron trajectory (**Fig. 2f, Extended Data Fig. 5e**). Previous studies have shown that degradation of neuronal identity is associated with increased glycolytic metabolism in organoids ^21,22^. Indeed, GO analysis of organoid DL neurons indicates dysregulation of neuron differentiation, regionalization, and pattern specification (**Fig. 2b**). To test whether glycolysis is associated with cell fate or cell identity anomalies in dup15q DL neurons, we explored the expression of well referenced DL and UL lineage markers in organoid L5_6 cells. We found that dup15q L5_6 cells had decreased expression of canonical DL markers (*FEZF2*, *NFIB*, *BCL11B*, and *NEUROD6*) together with increased expression of immature (*DCX* and *PTN*) and UL (*SATB2* and *MEF2C*) markers compared to controls (**Fig. 2g**). To validate this finding, we performed immunostaining of 100 DIV organoids and found an increased co-expression of deep and UL markers in dup15q L5_6 cells (**Fig. 2h**) not present in control neurons. Together, these findings suggest that increased glycolysis in dup15q DL neuronal lineages may lead to degraded layer identity.

## Spatially resolved transcriptomic analysis of dup15q tissue sections

To further explore cell-specific gene expression changes in dup15q with spatial resolution, we curated a list of 285 genes (31 cell identity markers and 254 differentially expressed genes) for spatial transcriptomic analysis (**Extended Data Table 5**). Using the MERSCOPE platform, we analyzed the spatial expression of targeted genes at sub-cellular resolution in the prefrontal cortex of dup15q and control samples. Pre-processed spatial data was piped into Seurat (v5) for dimensionality reduction, single-cell clustering, and cell annotation. The analysis identified excitatory neurons from all cortical layers, as well as subtypes of interneurons, glia, and mural cells (**Figs. 3a, b**). Using spatial coordinates, annotated clusters were plotted over the tissue image (**Fig. 3c**). Exploring the expression of layer-specific markers, despite overall lower RNA detection rates, we did not observe cellular organization anomalies in the dup15q sample (**Fig. 3c**, **Extended Data Fig. 6a**). However, we were able to validate many differentially expressed genes originally identified by snRNA-seq that were highly transcribed across multiple cell-types (**Extended Data Fig. 6b**), as well as cell-type specific changes at single cell resolution (**Fig. 3d**).

**Figure 3.**
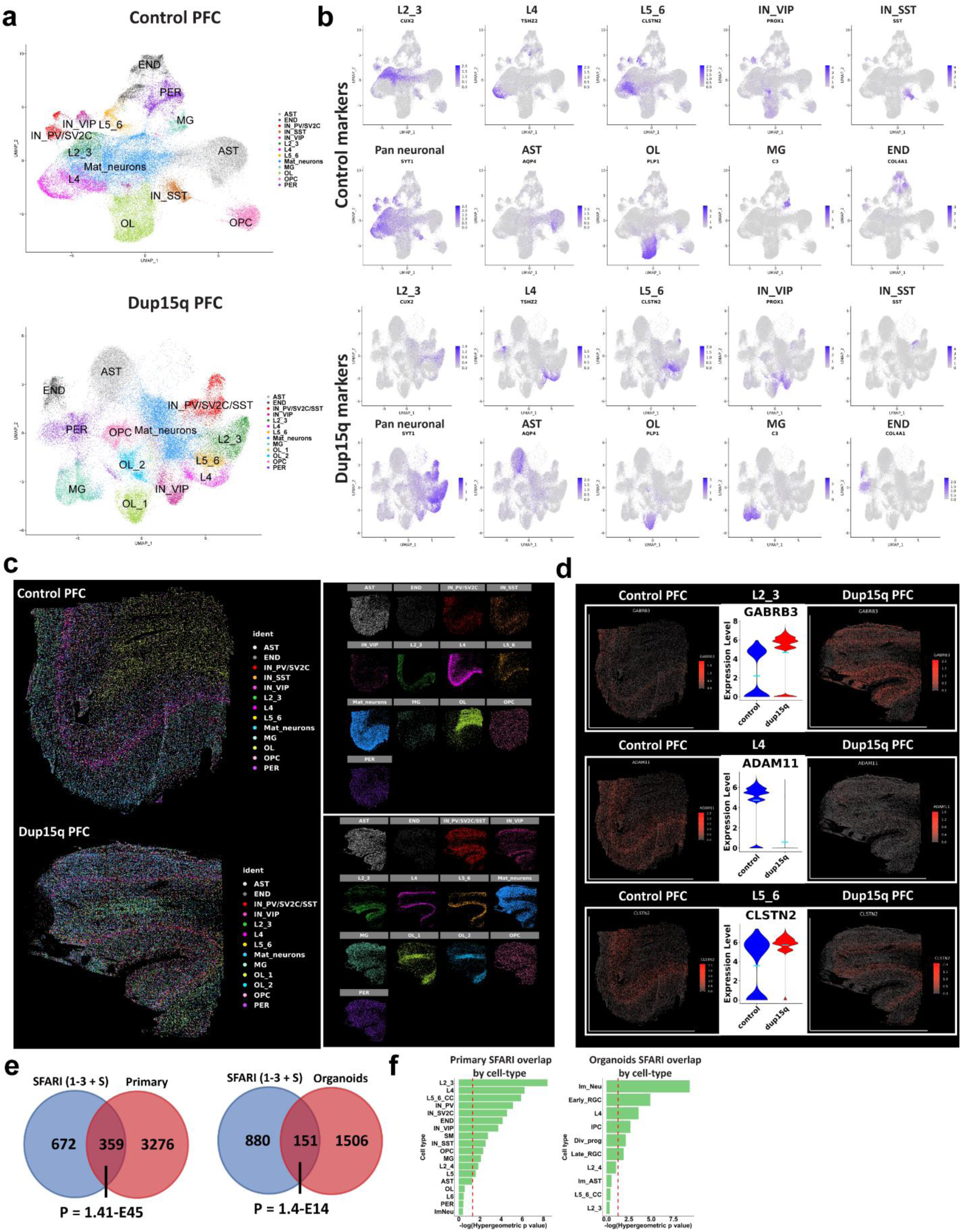
Spatial resolved transcriptomics of dup15q syndrome prefrontal cortex. **a)** Uniform Manifold Approximation and Projection (UMAP) embedding and cluster annotations of single cell spatial transcriptomic data. **b)** Cell subtype gene expression of markers used for cluster annotations. **c)** Annotated cell clusters overlaid on tissue images using cell coordinates. Image showing comparable cell type identification and spatial localization of dup15q patient and control clusters. **d)** Examples of cell type specific differential gene expression validated by spatial resolved transcriptomics. **e)** Overlap between primary and organoid DEGs and high-confidence ASD genetic risk factors (SFARI gene scores 1 to 3 and syndromic; Hypergeometric P-value). **f)** Overlap between SFARI genes and DEGs by cell-type; dashed red line indicates statistical significance (q < 0.05).

## Dup15q DEGs converge on known ASD risk genes

Hypothesizing that dup15q gene expression dysregulation coalesces to known ASD associated genes and pathways, we analyzed the intersection between all primary and organoid DEGs with the list of 1,031 high-risk ASD associated genes from the Simons Foundation Autism Research Initiative (SFARI) database ^37^ (rank 1-3 + syndromic, N = 1031; 1/11/22 release). Of 3,635 unique primary DEGs, 359 were found in the SFARI gene database (P = 1.41E-45, hypergeometric test). Of 1,657 unique organoid DEGs, 151 genes intersected with the SFARI dataset (P = 1.40E-14, hypergeometric test, **Fig. 3e**). This suggests that dup15q gene dysregulation is implicated in known ASD genetic pathways **(Extended Data Tables 2,3)**. Next, we sought to analyze this intersection with SFARI genes in specific cell types. In primary dup15q tissue, we found that SFARI genes were enriched in DEGs of excitatory neurons, particularly in UL neurons (L2_3), suggesting that the dup15q ASD pathophysiology is preferentially associated with DEGs in these cell-types. In organoids, immature neurons and early radial glia had the highest overrepresentation of SFARI genes **(Fig. 3f, Extended Data Tables 2,3)**. Interestingly, in contrast to postmortem ASD tissue, organoid L2_3 cortical neurons had the least number of SFARI overlapping genes. This might reflect the relatively immature transcriptomic state of UL neurons from organoids, which correspond to fetal stages, compared to primary nuclei from patients over 8 years of age ^22^.

## Convergence of synaptic gene dysregulation between dup15q and idiopathic ASD

We next sought to explore the potential convergence of cell specific molecular changes between dup15q and idiopathic autism spectrum disorder (ASD). Initially, we computed the Pearson’s correlation coefficient for highly expressed genes between corresponding clusters of the dup15q postmortem samples and our previously published ASD snRNAseq datasets ^8^. We found a high degree of gene expression correlation between corresponding clusters, showing that annotated cells from each of the datasets are indeed comparable **(Extended Data Fig. 6c)**. Remarkably, when we compared gene burden analyses of ASD and dup15q, we found that UL projection neurons were preferentially affected in both (**Fig. 2a**) ^8^. GO analysis of ASD and dup15q UL (L2_3) neuron DEGs showed enrichment for convergent pathways involving regulation of neuron projection and synaptic related terms such as synapse assembly, synaptic plasticity, synaptic organization, and synaptic transmission (**Fig. 4a**). When we examined the number of overlapping DEGs between UL neurons of dup15q and ASD, we found 24 overlapping genes (P = 3.15E-11, hypergeometric test) with 23/24 genes showing the same expression directionality, and a significant degree of gene expression correlation (Pearson’s R coefficient = 0.91, P = 1.1E-09; **Fig. 4b**). Overlapping genes were found to be associated with synaptic function and plasticity (*CNTNAP2*, *SLITRK5*, *CABP1*, *MAPK1* and *NFIA*) and with dendritic spine morphology (*CNTNAP2* and *CABP1*) (**Fig 4c**). Importantly, both neuron projection and synaptic function related terms have been consistently reported to be affected in postnatal ASD brains, highlighting conserved cell type specific molecular changes in UL projection neurons between dup15q and other ASD subtypes ^5,6,8^.

**Figure 4.**
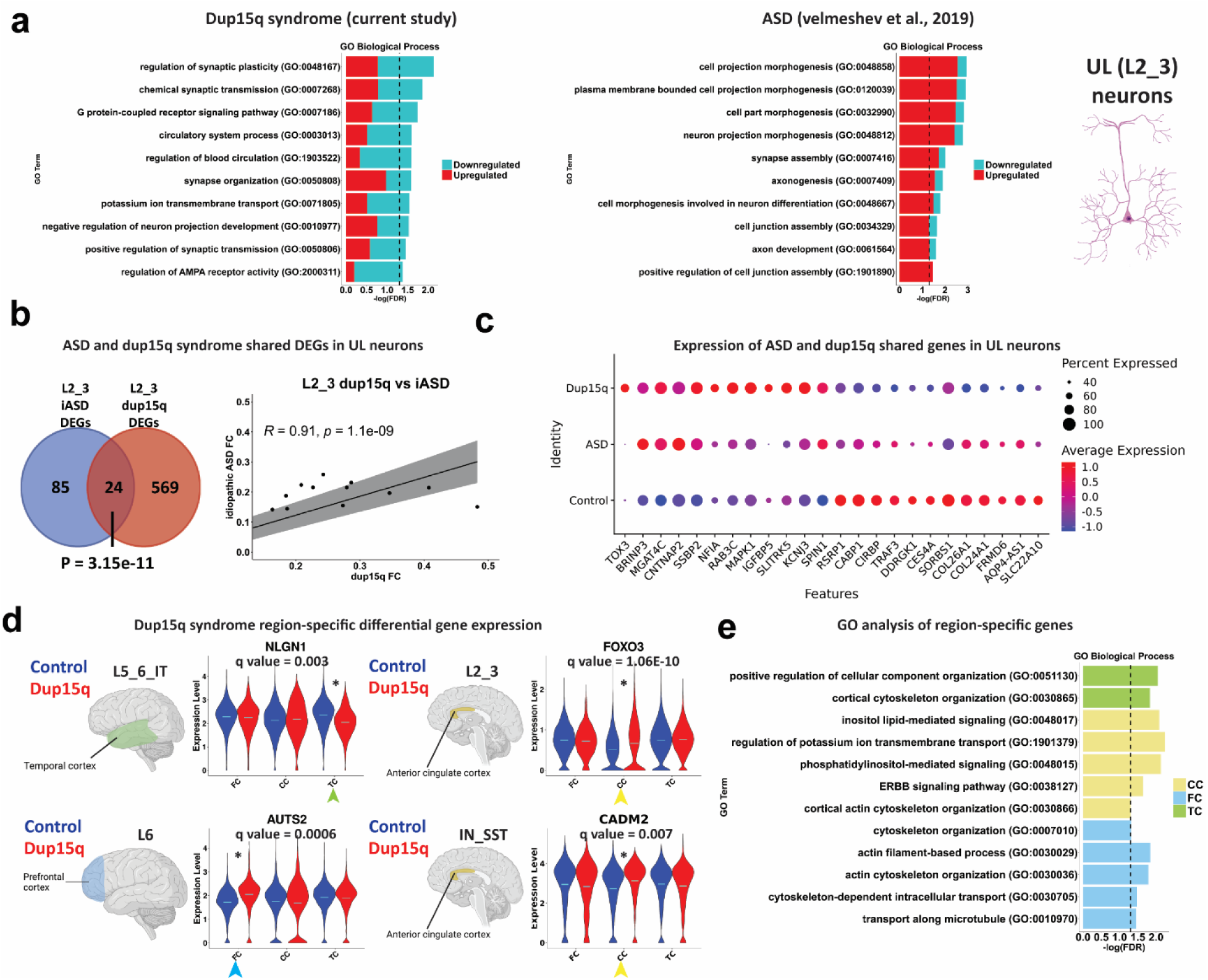
Converged molecular changes of UL neurons between dup15q syndrome and idiopathic ASD. **a)** GO analysis of idiopathic autism (ASD) and dup15q syndrome UL neuron DEGs. Overlap between upper-layer DEGs of idiopathic ASD and dup15q as well as Pearson’s correlation coefficient of shared gene fold changes. **c)** Differential expression of ASD and dup15q syndrome shared genes in UL cortical neurons. **d)** Violin plots of selected dup15q region-specific gene expression. **e**) GO analysis of dup15q region-specific genes.

## Cortical region-specific gene expression changes in dup15q

We next explored dup15q gene expression changes across different brain regions in a cell type specific manner (q value < 0.05; expression fold change ≥ 10%). We found 472 genes across all cell types that were differentially expressed between dup15q and controls in a region-specific manner. The majority of region-specific DEGs were found in the PFC (262 genes; 55.5%), followed by the TC (111 genes; 23.5%) and ACC (99 genes; 21.0%). Most region-specific DEGs occurred in excitatory neurons (230 genes, 48.7% of all region-specific genes), reflecting spatial heterogeneity of dup15q molecular changes within excitatory subtypes. Only 11 region-specific genes were strong enough to drive the overall cell type specific DEGs across all regions (6 in neurons and 5 in non-neuronal cells) **(Extended Data Table 6)**. Some of the cell type region-specific genes that we found were known ASD-associated genes (**Fig. 4d**). For example, in L5_6_IT neurons, *NLGN1* was downregulated in the temporal cortex of dup15q, but not in the PFC or the ACC. This ASD-associated gene encodes a cell-adhesion protein known for its role in synapse formation and long-term potentiation ^38^. In L6 corticofugal projection neurons, *AUTS2* was upregulated in the cingulate cortex (CC). This known ASD risk gene encodes a component of a polycomb group (PcG) multiprotein PRC1-like complex that is important for axon and dendrite elongation through regulation of neuronal gene expression during brain development ^39^. *CADM2*, another ASD associated gene that encodes an adhesion molecule important for synapse organization, was enriched in somatostatin interneurons (IN_SST) of the cingulate cortex of dup15q patient samples. We also found that *FOXO3*, an important determinant of neuronal morphogenesis and neuronal survival ^40,41^, was enriched in dup15q L2_3 projection neurons of the ACC. Gene ontology (GO) analysis showed enrichment for cortical cytoskeletal organization in all cortical regions (**Fig. 4e**). This might suggest that dup15q region-specific gene expression is implicated in cellular dysorganization or cytoarchitectural anomalies in these cortical regions. A previous publication showed neuronal soma volume deficits in the limbic system, and in other sub-cortical regions of dup15q patients ^42^. However, our spatially resolved transcriptomic analysis of dup15q, failed to identify such anomalies in the PFC.

## Gene co-expression analysis highlights autism associated developmental and postmortem modules

To further explore specific pathways that are dysregulated in dup15q syndrome, we performed high-dimensional weighted gene co-expression network analysis (hdWGCNA) on all cells of both dup15q postmortem and organoid datasets. We focused on excitatory intratelencephalic (IT) neurons in postmortem samples and organoids since they shared a high degree of gene expression burden. Co-expression network analysis identified 18 modules in organoids and 21 modules in postmortem samples (**Fig. 5a, Extended Data Tables 7,8**). While Primary tissue modules were variably expressed across all IT neurons, organoid modules were mostly enriched in L5_6 and L4 cells. (**Fig. 5b**). We then performed module trait correlation to focus on modules associated with dup15q. We highlighted two downregulated and two upregulated modules in each dataset that show a high degree of disease trait correlation and a relatively low correlation with age (primary) or stage of *in vitro* development (organoids) across all cell types (**Fig. 5c**). Modules IT2 and IT6 were downregulated and modules IT9 and IT10 were upregulated in primary nuclei of dup15q patients. GO analysis showed that primary IT neuron downregulated modules were associated with sphingolipid metabolism, axon guidance and neurotransmitter associated channel activity, whereas upregulated modules were associated with post-transcriptional mRNA processing, ion channel activity and post synaptic regulation and function (**Fig. 5d**). Amongst downregulated modules, we observed known autism risk genes associated with modulation of chemical synaptic transmission and synaptic plasticity such as the voltage-gated ion channels *CACNA1C*, *CACNA1D*, *CACNA1G, KCND3* and *KCNB1*, the ankyrin repeat containing family members *SHANK1*, *SHANK2* and *ANKRD11,* and the *GRIN2A* receptor subunit of the N-methyl-D-aspartate (NMDA) glutamate receptor. IT upregulated modules also included many genes involved in synaptic activity and localization of receptors to synapses such as the neuronal cell surface proteins, *NLGN1* and *NRXN1*, a regulator of synaptogenesis, *ADGRB3,* the synaptic spine architecture and function regulator, *LRRC7*, and *DLG1*, a scaffolding protein that is involved in recycling and clustering of glutamate receptors (**Fig. 5e, Extended Data Table 7**). Interestingly, studies of dup15q mouse models as well as from patient stem cell-derived neurons, demonstrate synaptic impairments leading to hyperexcitability ^43–45^. Our results, specifically involving the IT9 module may point to a specific cluster of genes acting together to contribute to glutamatergic neuron hyperexcitability and seizure phenotypes. In organoid glutamatergic neurons, modules Ex6 and Ex12 were downregulated and modules Ex9 and Ex10 were upregulated. GO analysis showed that the downregulated modules were mostly associated with microtubule dynamics and mRNA processing related to transcription regulation and splicing, whereas the upregulated modules were associated with neurotrophic stimulus response, synaptic transmission, and glycolytic metabolic processes (**Fig. 5f, Extended Data Table 8**). The latter supports our previous observation of glycolysis enrichment in dup15q progenitors and DL neurons. Remarkably, within the Ex6 downregulated module were many prominent ASD risk genes (SFARI gene score 1 and/or S) that are strongly involved in transcription regulation via chromatin remodeling such as *KMT2A*, *KMT2C*, *KMT2E*, *KMT5B, ASH1L* and *SETD5 (*all act as methyl-transferases), *CHD2* (chromodomain containing chromatin remodeler), *KANSL1* (subunit of histone acetylation complex), *KDM6B* (Histone demethylase), *POGZ* (zinc finger protein containing a transposase domain), *MBD5* (methyl-CpG-binding), *ARID1B* (component of the SWI/SNF chromatin remodeling complex), and *WAC* (a linker between gene transcription and histone H2B mono-ubiquitination) **(Fig. 5g, Extended Data Table 8)**. These results reveal temporal changes of co-expressed gene networks in dup15q glutamatergic neurons, highlighting cell type-specific enrichment of modules containing ASD risk genes.

**Figure 5.**
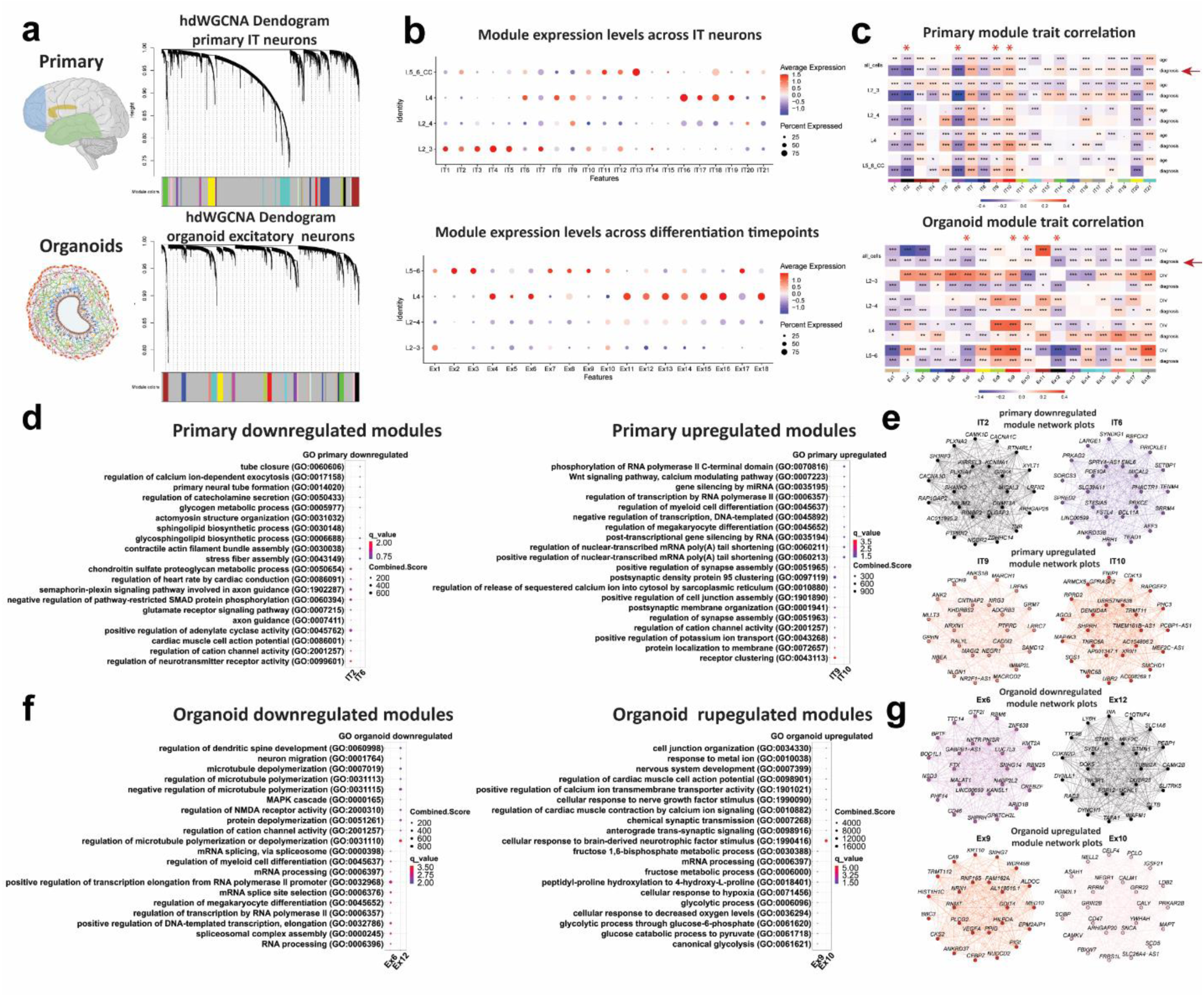
Weighted gene co-expression networks (WGCNA) of organoid and primary dup15q excitatory neurons. **a)** WGCNA dendrogram of primary IT neurons and organoid excitatory neurons. Each leaf represents a single gene, colors on the bottom represent assignment of co-expression modules. **b)** Primary IT neurons and organoid excitatory module expression across different cell-types. **c)** Module trait correlation analysis. Red Asterix indicates down or upregulated modules that are associated with the dup15q genotype. **d)** primary IT neuron gene ontology (GO) of down and upregulated dup15q associated modules. **e)** Individual module network plots of primary IT neurons highlighting the top 25 hub genes for each network. **f)** Organoid excitatory neuron gene ontology (GO) of down and upregulated dup15q associated modules. **g)** Individual module network plots of organoid excitatory neurons highlighting the top 25 hub genes for each network.

Numerous studies highlight gene expression regulation via chromatin remodeling and synaptic function as key mechanisms that drive ASD pathology ^2,5,7,46,47^. Our study provides further evidence that these distinct molecular pathways are subject to temporal molecular dysregulation in ASD brains. Dup15q associated organoid gene co-expression modules suggest that *in-utero* developmental pathologies are associated with neuronal transcription regulation and chromatin remodeling. These early molecular changes are likely driving synaptic maturation, plasticity, and neuronal circuit dysregulation during postnatal stages. This also suggests that addition of other brain regions and sampling time points might further uncover points of convergence for ASD ^11^.

## Conclusion

We identify cell-type specific dysregulated gene programs that underlie dup15q pathogenesis and show vulnerability of excitatory cortical neuron subtypes throughout development. Importantly, we show that dup15q is associated with dysregulation of neuron projection, especially in DL excitatory neurons, that is maintained at postnatal stages. Interestingly, we also find that metabolic dysfunction, namely an increase in glycolysis of dup15q progenitors, can lead to degraded neuronal identity of organoid DL neurons. We show that UL neurons are the most affected cell type postnatally, emphasizing that dup15q associated transcriptional burden is dynamically changing and might be correlated with maturation of specific cell types. While we could not find evidence for cellular organization anomalies in the PFC, we were able to validate differentially expressed dup15q syndrome genes at single cell resolution using spatially resolved transcriptomics. Importantly, we identify cortical region-specific gene changes that are associated with cytoskeletal organization and highlight brain region-specific changes in dup15q. Furthermore, we find convergent cell type-specific gene expression changes in both dup15q and idiopathic ASD, highlighting changes in synaptic regulation of UL neurons that are shared between syndromic and idiopathic ASD. This might indicate that these molecular changes are more relevant to the autism endophenotype of dup15q. Weighted gene co-expression analysis revealed temporal changes in gene networks expressed by dup15q glutamatergic neurons that are enriched for ASD risk genes. These findings highlight dynamic temporal shifts in the cellular and molecular impact of dup15q during neocortical development. Our data, describing transcriptomic changes across cortical cell types in dup15q can be viewed and interrogated via an interactive web browser: https://dup15q-cortex-organoids.cells.ucsc.edu.

## Acknowledgments

We thank Dr. Stormy Chamberlain and the University of Connecticut stem cell core for providing three dup15q and two of the control induced pluripotent stem cell lines used for this study.

## Author contributions

YP designed the project, assisted in acquiring funding, collected tissue, performed nuclei isolation and 10x Genomics capture, performed organoid differentiation and single cell capture, analyzed the data, performed spatial transcriptomic experiments, analyzed spatial transcriptomic data, and wrote the manuscript. DV designed the project, assisted in acquiring funding, collected tissue, performed nuclei isolation and 10x Genomics capture, assisted in data analysis, and edited the manuscript. LW assisted with hdWGCNA analysis. MW, JB and NGD performed organoid immunostainings. CS performed RNA in-situ hybridizations. SW performed tissue sections. MH created an interactive data visualization website. AK designed the project, acquired funding, supervised the project, and edited the manuscript. All authors read the manuscript.

## Data availability

Raw data can be accessed at the Sequence Read Archive (SRA), accession numberPRJNA1017130. Analyzed data (cell-count matrix and metadata) can be accessed through the UCSC Cell Browser, collection dup15q-cortexl-organoids (https://dup15q-cortex-organoids.cells.ucsc.edu).

## Supplementary Materials

**Extended Data Table 1. Sample metadata.**

**Extended Data Table 2. Nuclei-specific differentially expressed genes of dup15q cortex.**

**Extended Data Table 3. Cell-type specific differentially expressed genes of dup15q organoids.**

**Extended Data Table 4. Lineage-specific differentially expressed genes of dup15q organoids.**

**Extended Data Table 5. Genes used for spatial transcriptomics.**

**Extended Data Table 6. Cortical region-specific differentially expressed genes.**

**Extended Data Table 7. Weighted gene co-expression modules of cortical IT neurons.**

**Extended Data Table 8. Weighted gene co-expression modules of organoid excitatory neurons.**

### Materials and Methods

#### Cortical tissue sample preparation

Dup15q snap-frozen post-mortem tissue and control samples were obtained from the Autism BrainNet bank. Additional control samples were collected from the NIH NeuroBioBank. Cortical samples were sectioned using a cryostat, across the entire cortical span (as much as possible) to include grey matter and adjacent white matter. Sections were initially collected (∼10mg) for RNA extraction following an RNA integrity (RIN) assay using 2100 Bioanalyzer and an RNA pico Chip (Agilent). Larger sections (∼40mg) of high quality (RIN>6.9, unless they were dup15q samples) were collected for nuclei extraction.

#### Stem cell characterization maintenance and cortical organoid differentiation

In this work we used the following human induced pluripotent stem cell lines (hiPSC): Dup1-8, Rx68i, SCC115, UCH-CBY4-48B and UCH-PB-48E, obtained and authenticated by the University of Connecticut stem cell core. We also used the 13234 line from the Conklin lab (Gladstone Institute). All stem cell lines used in this study undergone G-banding karyotyping to validate either the dup15q or control genotypes. All lines were karyotyped and analyzed in WiCell (Madison, WI). All control lines showed normal karyotype without any genomic abnormalities or mosaicism. All dup15q lines reports identified the expected abnormality (idic(15q) karyotype for DUP1-8 and Rx68i and interstitial triplication for SCC115).

All stem cell lines were validated for pluripotency after receipt. Every 10 passages, stem cells were tested for karyotypic abnormalities and validated for pluripotency markers Sox2, Nanog, and Oct4. All cell lines tested negative for mycoplasma.

All hiPSC lines were expanded on growth factor-reduced Matrigel (BD)-coated plates. hiPSCs were thawed in StemFlex Pro Media (Gibco) supplemented with 10 µM Rock inhibitor Y-27632 which was removed the following day. Medium was changed every day and lines passaged when colonies reached 80% confluency. Stem cells were passaged using ReLeSR (Stemcell technologies) and manually lifted with cell lifters (Fisher). All lines used for this study were bellow passage 30.

Cortical organoids were cultured using a dorsal forebrain differentiation protocol ^20^. Briefly, hiPSC lines were expanded and dissociated to single cells using accutase (Gibco). Cells were then aggregated in neural induction medium at a density of 10,000 cells per well in 96 well v-bottom low adhesion plates. GMEM-based neural induction medium included 20% Knockout Serum Replacer (KSR), 1X non-essential amino acids, 0.11 mg/mL Sodium Pyruvate, 1X Penicillin-Streptomycin, 1X Primocin, 0.1 mM Beta Mercaptoethanol, 5uM SB431542 and 3 µM IWR1-endo. Medium was supplemented with 20 µM Rock inhibitor Y-27632 for the first week. After 18 days organoids were transferred to six well ultra-low adhesion plates and moved to an orbital shaker rotating at 90 rpm. Media was changed to DMEM/F12-based medium containing 1X Glutamax, 1X N2, 1X CD Lipid Concentrate and 1X Penicillin-Streptomycin. At 35 days, organoids were moved into DMEM/F12-based medium containing 10% FBS, 5 µg/mL Heparin, 1X N2, 1X CD Lipid Concentrate and 0.5% Matrigel. At 70 days medium was additionally supplemented with 1X B27 and Matrigel concentration increased to 1%. Organoids were collected for single cell capture and immunohistochemistry at relevant timepoints.

#### Nuclei isolation and snRNA-seq

Dup15q and matched control samples were processed as previously described ^48^. Briefly, sectioned brain tissue was homogenized in 5 mL of RNAase-free lysis buffer (0.32M sucrose, 5 mM CaCl_2_, 3 mM MgAc_2_, 0.1 mM EDTA, 10 mM Tris-HCl, 1 mM DTT, 0.1% Triton X-100 in DEPC-treated water) using a glass dounce tissue homogenizer (Thomas Scientific, Cat # 3431D76) on ice. The homogenate was loaded into a 30 mL thick polycarbonate ultracentrifuge tube (Beckman Coulter, Cat # 355631). 9 mL of sucrose solution (REF) (1.8 M sucrose, 3 mM MgAc_2_, 1 mM DTT, 10 mM Tris-HCl in DEPC-treated water) was added to the bottom of the tube with the homogenate and centrifuged at 107,000 g for 2.5 hours at 4 C. Supernatant was aspirated, and nuclei pellet was incubated in 250 uL of DEPC-treated water-based PBS for 20 min on ice before resuspending the pellet. Nuclei were filtered 3 times to remove debris using a 30*μ*m pre-separation filters (Miltenyibiotec). Nuclei were counted using a hemocytometer and diluted to 1,000 nuclei/uL before performing single-nuclei capture on the 10X Genomics Single-Cell 3’ system. Target capture of 10,000 nuclei per sample was used. Nuclei capture and library preparation was done according to the Chromium Single Cell 3’ Reagent Kits User Guide V3 (10x genomics). Single-nuclei libraries were pulled and sequenced as per manufacturer recommendation on a NovaSeq S2 flow cell.

#### Cortical organoid single cell capture and scRNA-seq

Organoids were collected for single cell capture and immunohistochemistry (N=3 organoids per line per timepoint) at 50, 100 and 150 days of *in-vitro* development (DIV). Organoids were dissociated using papain (Worthington) solution containing DNase. Samples washes 3 times in 1XPBS, placed in 1.5 ml papain and incubated at 37 °C. tubes were inverted every 15 minutes for up to a total of 45 minutes. Next, samples were triturated by manually pipetting with a glass pasteur pipette approximately until homogenized. Dissociated cells were spun down at 300g for 5 min and papain removed. Similar to nuclei capture, 10,000 cells per sample was used and library preparation was done according to the Chromium Single Cell 3’ Reagent Kits User Guide V3 (10x genomics). Single-cell libraries were pulled and sequenced on a NovaSeq S2 flow cell.

#### 10X Genomics CellRanger software and data filtering

CellRanger software v 6.1.1 was used for library demultiplexing, fastq file generation, read alignment and UMI quantification. CellRanger was used with default parameters using the GRCh38-2020-A transcriptome reference, while using the “include-introns” option as pre-mRNA reference file for nuclei data. These were piped into Seurat v.4. For nuclei analysis ^49,50^, expression matrices containing numbers of Unique molecular identifiers (UMIs) per nucleus per gene were filtered to retain nuclei with at least 500 genes expressed and less than 5% of total UMIs originating from mitochondrial and ribosomal RNAs (nCount_RNA > 500, percent.mt < 5% and percent.ribo < 5%). Individual matrices were combined, UMIs were normalized to the total UMIs per nucleus and log transformed.

For organoid cell analysis expression matrices containing numbers of UMIs per cell per gene were filtered to retain cells with at least 1000 genes expressed and less than 10% of total UMIs originating from mitochondrial and 25% ribosomal RNAs (nCount_RNA > 1000, percent.mt < 10% and percent.ribo < 25%).

#### Dimensionality reduction, clustering, cell type annotation and UMAP visualization

Data preprocessing, normalization, variable feature selection and PCA was performed using the standard Seurat (v.4.) pipeline ^49,50^. Briefly, filtered log-transformed UMI matrix was used to perform principal component analysis. Scree plot was generated to select the number of significant principal components (PCs) by localizing the last PC before the explained variance reaches plateau. Using this approach, k = 10 nearest neighbours were used for both postmortem nuclei and organoid cells to calculate Jaccard distance-weighted nearest neighbor distances which was used to perform Louvain clustering. To visualize transcriptomic profiles in two-dimensional space, we used Uniform Manifold Approximation and Projection for Dimension Reduction (UMAP) technique. The identity of specific cell types was determined based on expression of known marker genes, as shown in Figure 1 and figure S2c-d. One primary nuclei cluster that was previously identified as neurogranin (NRGN) expressing neurons ^48^ was annotated as neuronal debris due to nuclear-retained transcript depletion and high content of mitochondrial gene expression ^51^ (**fig S2e-f**).

#### Differential gene expression analysis

We downsampled the data so that the control and dup15q groups across all cell types had the same number of nuclei/cells. To identify genes differentially expressed in dup15q samples for both primary nuclei and organoid cell-types we used MAST to perform zero-inflated regression analysis by fitting a linear mixed model (LMM) ^52^. To exclude gene expression changes stemming from confounders, such as age, sex, RIN, cortical region, 10X capture batch and sequencing batch, the following model was fit with MAST for primary nuclei:

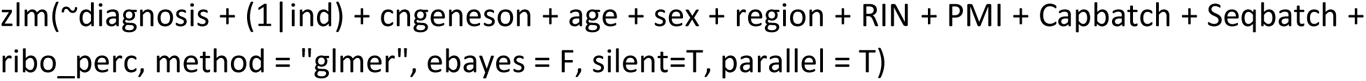

and for organoid cells:

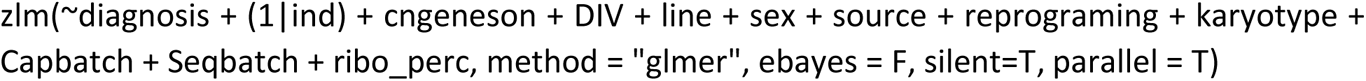

Where cngeneson is gene detection rate (factor recommended in MAST tutorial), Capbatch is 10X capture batch, Seqbatch is sequencing batch, ind is individual label, PMI is postmortem interval, DIV is days of *in-vitro* differentiation, source is the source tissue cells were reprogrammed from, reprograming is the method used for iPSC generation, ribo_perc is ribosomal RNA fraction, mito_perc is mitochondrial RNA fraction and mito_nucl_perc is the fraction of nuclear-encoded mitochondrial RNA.

When testing for differential expression in each cluster, the following strategy was used to prevent individual bias from affecting differential gene expression analysis. First, average number of cells contributed by each individual sample to a given cluster was calculated. Then, individuals contributing less than 25% of this average were removed from the analysis. Equal number of cells was selected from each sample, and differential expression was performed. This routine was repeated 10 times to sample most cells in each cluster. On average across clusters, this strategy allowed including 85% of all individuals when performing the analysis while preventing nuclei profiles from few individuals from dominating differential gene expression estimates.

In addition, we calculated raw fold change of gene expression by repeating MAST analysis with only diagnosis factor in the model and filtered out genes with raw fold change of expression less than 0.1. The latter filtering step allowed removing genes whose fold change of expression was heavily dependent on the confounding factors, rather than clinical diagnosis.

#### Permutation analysis

To verify the ability of our model to capture true ASD-associated gene expression changes, we preformed random permutation of individual labels in each cluster between control and dup15q groups before performing differential gene expression analysis. This analysis was repeated over 10 permutations.

#### Burden analysis

To calculate number of dup15q-associated DEGs across cell types and normalize by number of cells in each cluster, we randomly drew 500 nuclei/cells from the control and ASD groups and each cell type before performing differential expression analysis. This analysis was repeated across 10 permutations, and average number of DEGs for each cell type was estimated.

#### Trajectory reconstruction and identification of lineage-specific dynamically expressed genes in the organoid dataset

Seurat UMAP coordinates were imported into monocle3 ^53^ for trajectory reconstruction. learn_graph function with custom graph_control options were used to construct the trajectory graph. To focus on specific lineages, we isolated the shortest path between the node in the neural progenitor/radial glia cluster and the node in the more mature cell type clusters. Then, we selected cells along the trajectory branches corresponding to specific lineages. After that, monocle3’s Moran’s test (graph_test function) was used to identify genes that are dynamically expressed in each lineage in control and dup15q organoids separately. We modified graph_test function to utilize Moran’s test with covariates to ensure that our results are not affected by uneven contribution of cells from controls and Dup15q subjects, as well as cells from different iPS cell lines. We selected genes with adjusted p value < 0.05 and Moran’s I>=10 as statistically significant dynamically expressed genes. To identify lineage-specific genes, we first compressed the single-cell expression data along each lineage by using a sliding window along pseudotime and averaging expression of neighboring cells for each gene. We generated 500 meta-cells in each lineage using this approach.

#### Identification of dynamically expressed genes dysregulated in dup15q organoids in specific lineages

To identify dynamically expressed genes dysregulated in dup15q organoid lineages compared to controls, we selected all genes dynamically expressed in control and dup15q organoids in each lineage. Then, for each lineage and gene, we calculated the difference of the areas under the curve for the smoothed expression/pseudotime plot for each gene in each lineage between control and dup15q and termed it differential expression score. All genes with the differential expression score of at least 50 were considered differentially expressed between control and dup15q.

For plotting combined gene expression for all genes in specific pathways (glycolysis, oxidative phosphorylation, citric acid cycle), we used curated gene sets from MSigDB ^54^ (HALLMARK_GLYCOLYSIS, HALLMARK_OXIDATIVE_PHOSPHORYLATION and KEGG_CITRATE_CYCLE_TCA_CYCLE). We then calculated averaged gene expression of all genes in a pathway in each meta-cell and each lineage separately for control and dup15q organoids. Log-transformed values of averaged expression were plotted over pseudotime.

#### Region-specific differential gene expression analysis

To test for age and diagnosis interaction, the following LMM model was fitted:

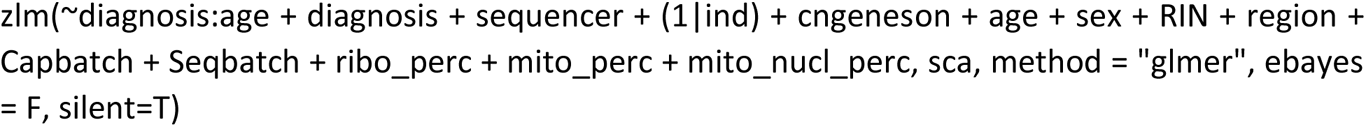

And LRT test was performed after removing diagnosis:age interaction term from the model. Genes with FDR<0.05 were considered as significantly affected by interaction between age and dup15q diagnosis. This analysis was performed for all clusters.

For region-specific analysis, nuclei from either the PFC, ACC or TC were selected, and the same LMM model was utilize as when testing for differential gene expression between Dup15q and Control, except for removing the region factor.

#### Spatially resolved transcriptomics

Sample preparation was performed according to manufacturer’s instructions (MERSCOPE Fresh and Fixed Frozen Tissue Sample Preparation User Guide, Doc. number 91600002). Briefly, fresh snap frozen tissues from the prefrontal cortex of dup15q syndrome patient and control, having a high RNA integrity number (RIN>8) were sectioned (10um thick) using a cryostat and mounted on MERSCOPE functional slides. Sections where then fixed and stored at 70% ethanol for up to two weeks. Sections went through autofluorescence quenching under UV light for 3 hours using the MERSCOPE Photobleacher instrument. A Pre-designed panel mix (285 genes) focused on highly differentially expressed genes based on the single-nuclei analysis were used for probe hybridization. Hybridizations were performed at 37°C for up to 48 hours in a humid environment. Post prob hybridization, sections were fixed using formamide and embedded in gel. After gel embedding, tissue samples were cleared using a clearing mix solution supplemented with proteinase K for 24-48 hours at 37°C until no visible tissue was evident in the gel. After clearing was completed, sections were stained for DAPI and PolyT and fixed with formamide prior to imaging. The MERSOPE imaging process was done according to the MERSCOPE Instrument Site Preparation Guide, Doc. Number 91500001. Briefly, an imaging kit was thawed at 37°C for 45 minutes, activated and loaded into the instrument. MERSCOPE flow chamber was then assembled with the stained tissue section, fluidics were primed, flow chamber filled with liquid and a low-resolution image was taken. Based on DAPI staining, an ROI was chosen for the full imaging experiment. After imaging was complete, data was processed using MERSCOPE proprietary software, while using DAPI and polyT staining for cell segmentation and RNA puncta cellular assignment. Further analysis, visualization, and integration of spatial data, was done using Seurat v5 (Source: vignettes/spatial_vignette_2.Rmd).

#### High-dimensional weighted gene co-expression network (hdWGCNA)

To construct hdWGCNA network from single nuclei/cell RNA-seq data, we used the hdWGCNA R package ^55^. initially, we subset the Seurat object to include IT excitatory neurons from nuclei and all excitatory cells from organoids. We then set up a Seurat object for WGCNA by selecting genes that are expressed in at least 5% of cells in each dataset. We continued by constructing metacells using k-Nearest Neighbors (KNN=25) to yield a metacell gene expression matrix. Following normalization and scaling of the data, we tested different softpowers to select for thresholding when performing WGCNA. When constructing co-expression network, we used soft_power=10 for primary IT neurons and soft_power=8 for organoids. We then computed all module eigengenes (MEs) in the full single-nuclie/cell datasets, harmonized module eigengenes and computed eigengene-based connectivity (kME) for each gene. We also performed module trait correlation to generate a heatmap of disease associated modules and used enrichR ^56^ to perform module specific gene ontology enrichment. Finally, we used ModuleNetworkPlot function to visualize the network underlying the top 25 hub genes for each disease associated module.

#### Dup15q and idiopathic ASD clusters gene expression correlation

To account for library size and number of cells per cluster, normalized average counts of each cluster from each dataset was obtained using Seurat’s AverageExpression function. For comparison, top 20 markers from each cell-type were used. Counts were converted to counts per million (CPM) and log transformed. Correlation analysis and plotting was performed using the corrplot R package.

#### Statistical overrepresentation test for Gene Ontology (GO) terms

PANTHER ^57^ was used to perform statistical overrepresentation test for DEGs from each cluster. All genes expressed in a given cluster were used as the background and GO Biological Processes ontology was used. Processes with FDR<0.05 were considered and sorted by FDR.

#### Hypergeometric testing

To estimate significance of overlap of two gene lists, we performed hypergeometric testing. To estimate overlap of ASD genetic risk factors with DEGs in each cell type, list of DEGs in a given cell type was used as the sample and list of all genes expressed in the cell type as the population list. These lists were overlapped with all genes in the SFARI Gene Module database, genes with ranks 1 to 3 or genes labeled as “syndromic”. Hypergeometric p values were FDR-corrected using Benjamini and Hochberg procedure.

#### Histology and immunohistochemistry

Human brain tissue blocks were snap-frozen and stored at −80°C. 16µm-cryosections were collected on superfrost slides (VWR) using a CM3050S cryostat (Leica) and fixed in 4% PFA at room temperature (RT). Organoids were captured at 50, 100 and 150 DIV, washed with 0.1M PBS and fixed in 4% PFA at room temperature (RT) for 1 hour. Organoids were then washed with 0.1M PBS and moved to a solution of 30% sucrose at 4°C overnight. Organoids were snap-frozen and stored at −80°C in a cryomold filled with a 1:1 30% sucrose to OCT ratio. For both organoid and brain tissue immunohistochemistry, 16µm sections were initially treated with heated (90°C) citrate-based antigen unmasking solution (Vector laboratories) and blocked in 0.1M PBS/0.1% Triton X-100/ 10% goat/horse/donkey serum for 30min at RT. Primary antibody incubations were carried out overnight at 4°C. After washing with 0.1M PBS, cryosections were incubated with secondary antibodies diluted in 0.1M PSB/ 0.1% Triton X-100 for 2 hours at RT. For immunofluorescence, Alexa fluochrome-tagged secondary IgG antibodies (1:500, Invitrogen) were used for primary antibody detection. Slides with fluorescent antibodies were mounted with DAPI Fluoromount-G (SouthernBiotech) and imaged on a Leica TCS SP5 confocal microscope. For intermixed neuron identity analysis, images were analyzed using Imaris x64 9.7.1. SATB2 (depicted in green) and BCL11B (depicted in red) were counted automatically. Overlapping was detected using the Imaris filter “shortest distance to” (lower threshold 0, upper threshold 1.5). Numbers shown are SATB2 and BCL11B double positive cells normalized by CTIP2 positive cells. For PGK1 expression, positive cells were quantified using the imaris spots detection function (minimum diameter threshold 7 and minimum quality filter threshold 11). The plot represents the percentage of PGK1 positive cells relative to DAPI.

#### Single-molecule *in situ* RNA hybridization

Tissue blocks were sectioned using a Cryostat (Leica) at 10 μm onto glass cover slides and stored at −80°C. For RNA in situ hybridization, the RNAscope Multiplex Fluorescent Reagent Kit v2 was purchased from ACD Advanced Cell Diagnostics and RNAscope probes against UBE3A (Hs-UBE3A; #570691), PLXNA4 (Hs-PLXNA4-C2; #460041-C2) and GABRG3 (Hs-GABRG3-C3; #525691-C3) were purchased from Advanced Cell Diagnostics (ACD biosciences). As fluorophores, Opal 520 (1:500), Opal 570 (1:750) and Opal 690 (1:750) (Akoya Biosciences) were used. For sample preparation, slides were fixed in prechilled 4% PFA for 1 hr at 4°C and rinsed twice with PBS. Sections were dehydrated in a sequential ethanol gradient of 50%, 70%, 100%, 100% EtOH for 5 min. at RT each. Sections were air dried for 5 mins and a hydrophobic barrier was drawn around each section and dried for 5 min at RT. 5 drops of RNAscope hydrogen peroxide were added to each section and incubated for 10 min at RT. Slides were washed twice with DI water. 5 drops of protease IV were added to each section and incubated for 30 mins at RT. Slides were washed twice with 1X PBS. RNAscope probes were hybridized for 2 hrs at 40°C and all subsequent steps were conducted according to the manufacturer’s recommendation (RNAscope Multiplex Fluorescent Reagent Kit v2 User Manual, #323100-USM/Rev Date: 02272019).

#### RNAscope Imaging, quantifications, and statistics

Initially, grey and white matter were distinguished using nuclei morphology and immunohistochemical staining for *NEUN* and *MBP*. Images for quantifications of RNA in situ hybridization were focused on regions of grey matter. Images were acquired using a confocal microscope (Leica TCS SP5 X, 63X objective). The investigator was blinded to sample labels before acquiring and processing the images, and all image processing steps were performed with the same parameters for all sections and individuals. Seven fields of view were taken for each sample. All fluorescent pictures are z-stacked. RNA counts were detected and quantified by Imaris software while using the same detection parameters for both control and dup15q images. RNA counts were then normalized to number of nuclei detected by DAPI for each image. For statistical analysis, an unpaired two-tailed student’s t-test was performed using the Graphpad prism 8.0 software.

#### Antibodies

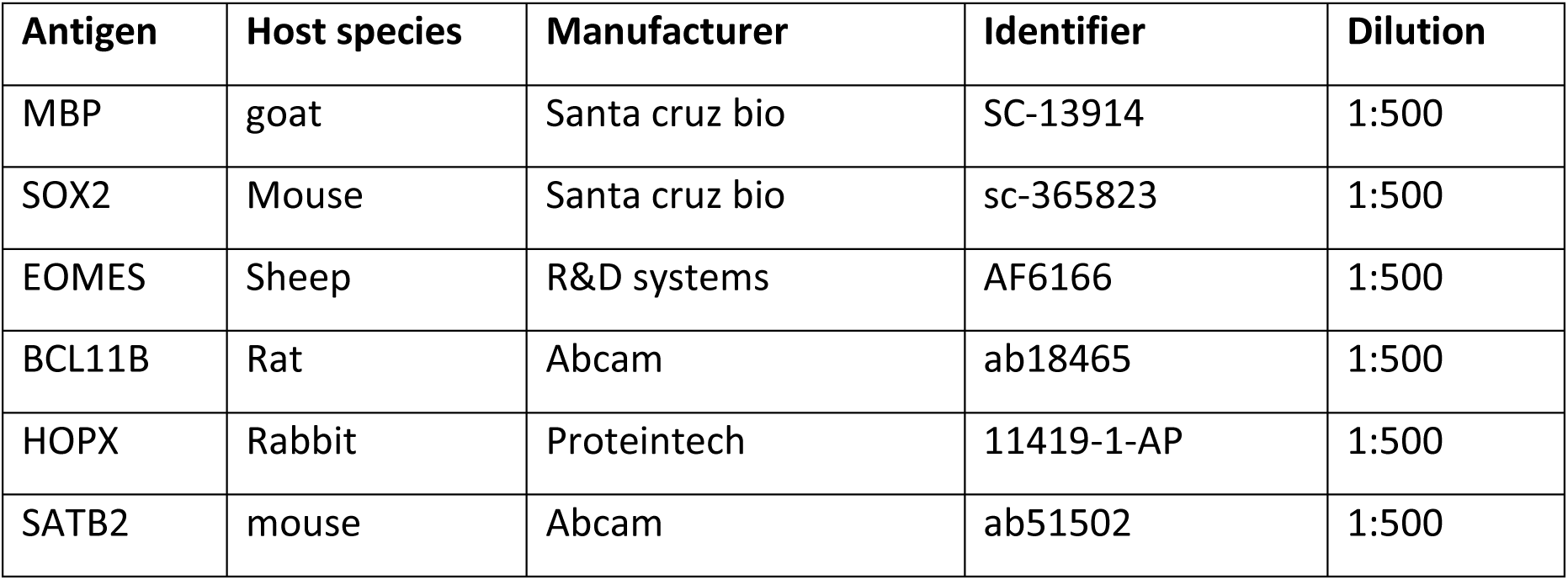
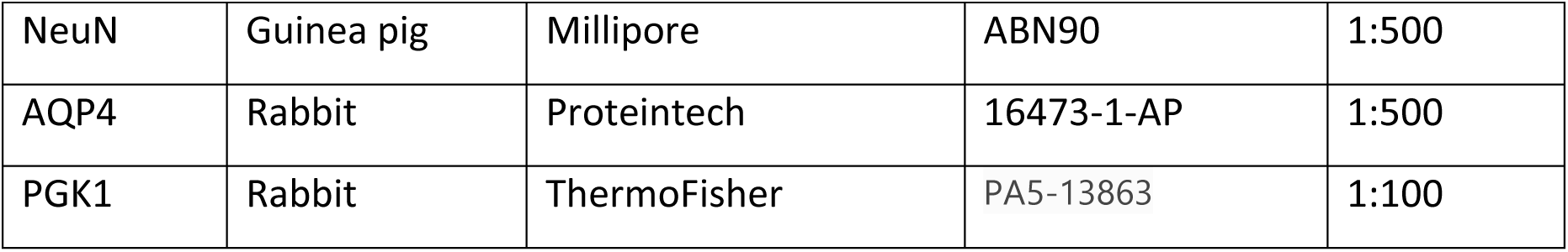

**Antibodies:**
| Antigen | Host species | Manufacturer | Identifier | Dilution |
| --- | --- | --- | --- | --- |
| MBP | goat | Santa cruz bio | SC-13914 | 1:500 |
| SOX2 | Mouse | Santa cruz bio | sc-365823 | 1:500 |
| EOMES | Sheep | R&D systems | AF6166 | 1:500 |
| BCL11B | Rat | Abcam | ab18465 | 1:500 |
| HOPX | Rabbit | Proteintech | 11419-1-AP | 1:500 |
| SATB2 | mouse | Abcam | ab51502 | 1:500 |
| NeuN | Guinea pig | Millipore | ABN90 | 1:500 |
| AQP4 | Rabbit | Proteintech | 16473-1-AP | 1:500 |
| PGK1 | Rabbit | ThermoFisher | PA5-13863 | 1:100 |

---

**Extended Data Figure 1.**
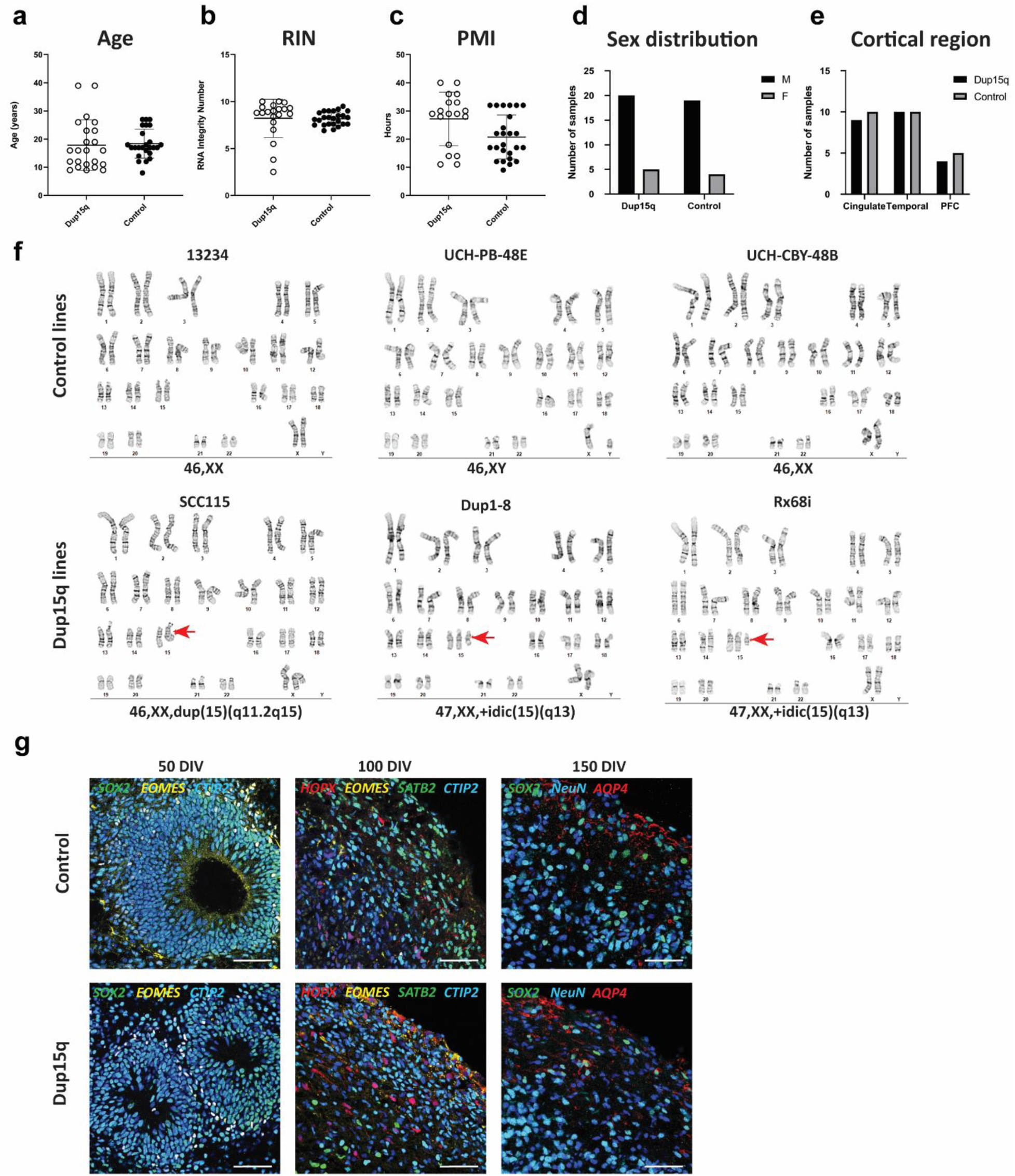
Sample statistics for experimental groups, iPSC karyotype and organoid development. **a-e)** Comparison of age, RNA integrity number (RIN), post-mortem interval (PMI), sex, and brain regions between control and dup15q groups. **f)** G-banding karyotype analysis of all iPSC lines used in the study**. g)** Immunostainings of organoid sections at three sampling developmental timepoints (50, 100 and 150 days of *in-vitro* development, scale bar = 50*μ*m).

**Extended Data Figure 2.**
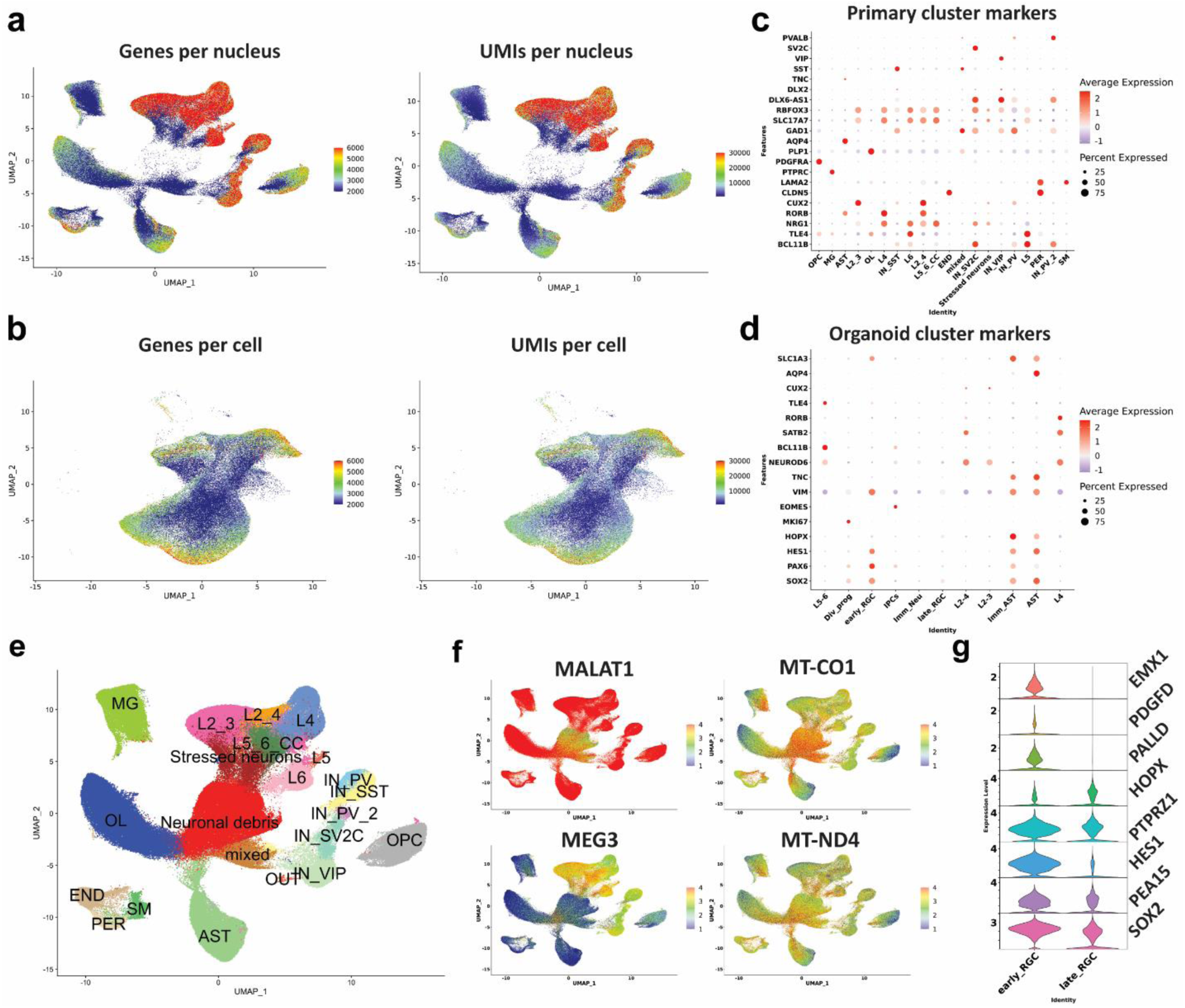
Technical and biological characteristics of the primary and organoid datasets. **a)** Primary gene and UMI counts per nucleus across all cell types. **b)** Organoid gene and UMI counts per cell across all cell types. **c)** Primary nuclei marker gene expression used to annotate specific cells. **d)** Organoid marker gene expression used to annotate specific cells. **e-f)** Identification of clusters containing neuronal debris, expression low nuclear-retention and high mitochondrial transcripts. **g)** Marker gene expression of early radial glia (early_RGC) vs late radial glia cells (late_RSC).

**Extended Data Figure 3.**
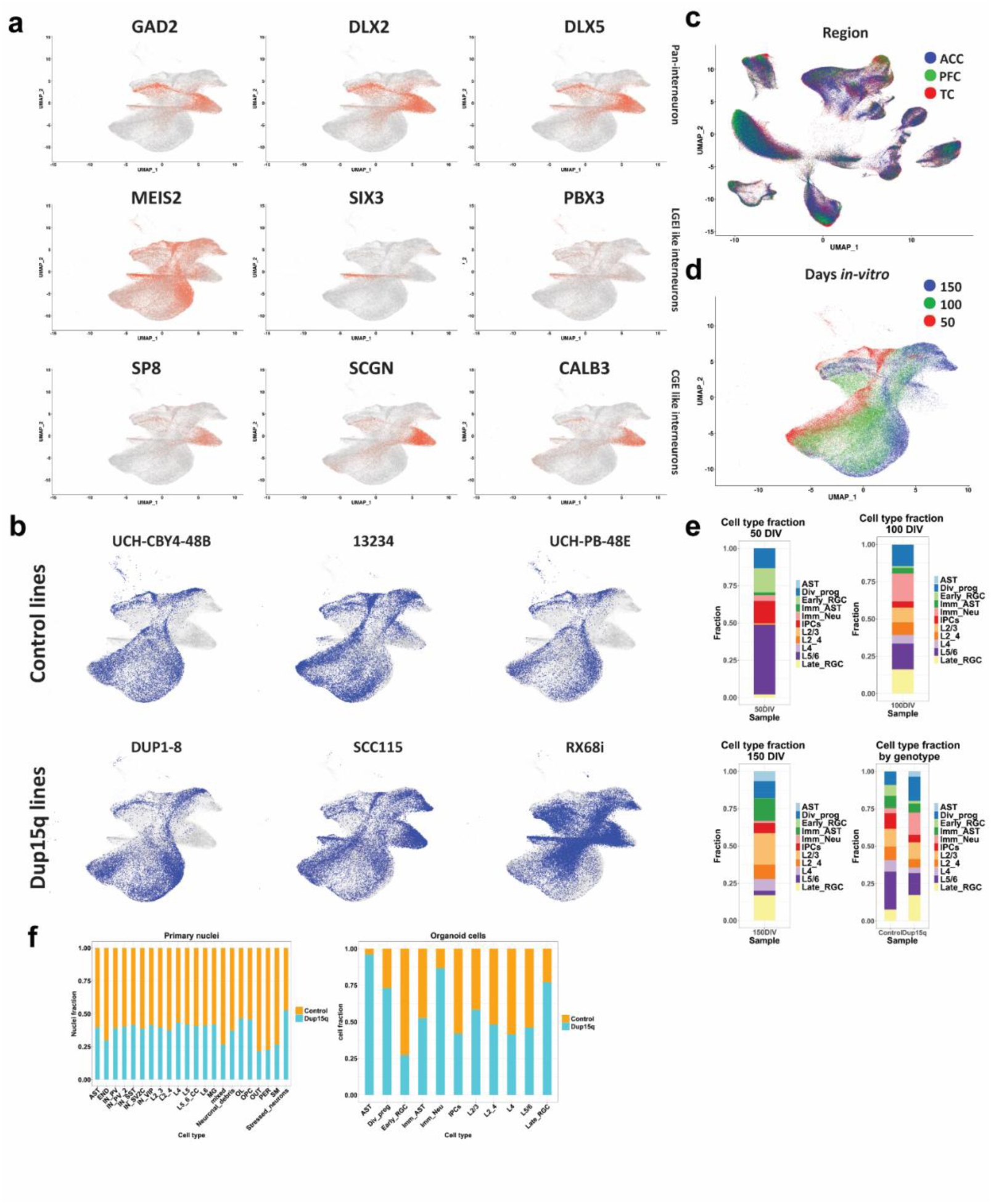
Organoid interneuron identification and cell-type proportions. **a)** Organoid marker gene expression identifies interneuron subtypes. **b)** Contribution of each line to specific organoid cell types. **c)** UMAP of all primary cell-types grouped by cortical region. **d)** UMAP of cell-types grouped by differentiation timepoint. **e)** Organoid cell type contribution for each differentiation timepoint, and between genotypes for all timepoints combined. **f)** Cell-type fraction from all primary nuclei and organoid cells type by genotype.

**Extended Data Figure 4.**
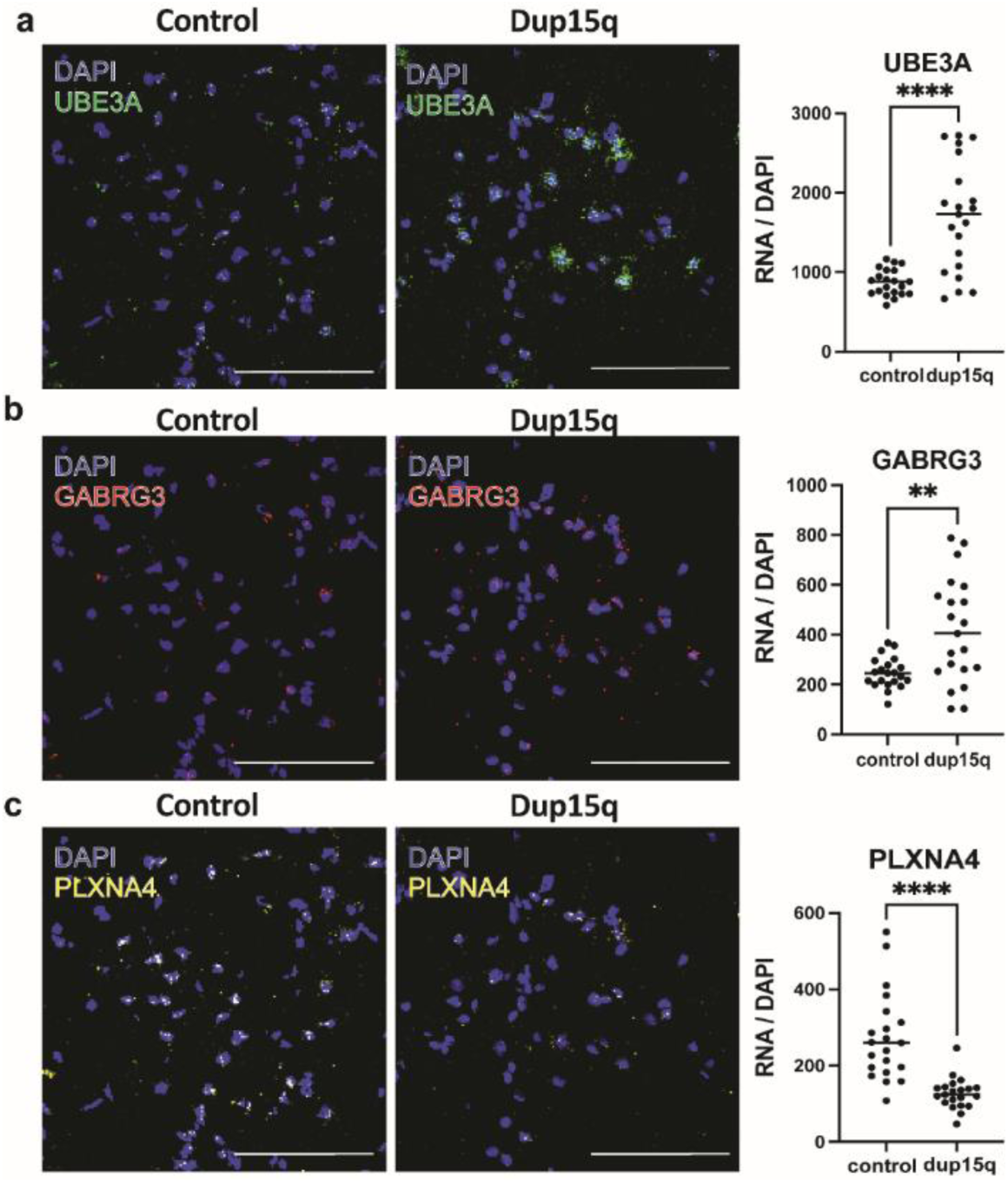
Single-molecule RNA *in situ* hybridization validation of selected differentially expressed genes across all cell types of dup15q and control cortical samples. **a)** Validation of enrichment of *UBE3A* transcript in neurons of the temporal cortex. **b)** Validation of enrichment of *GABRG3* transcript in neurons of the temporal cortex. **c)** Validation of *PLXNA4* transcript downregulation in neurons of the temporal cortex. (n=3 cortical samples from both dup15q and controls. Statistics represents counts from 7 field. Scale bar = 100µm).

**Extended Data Figure 5.**
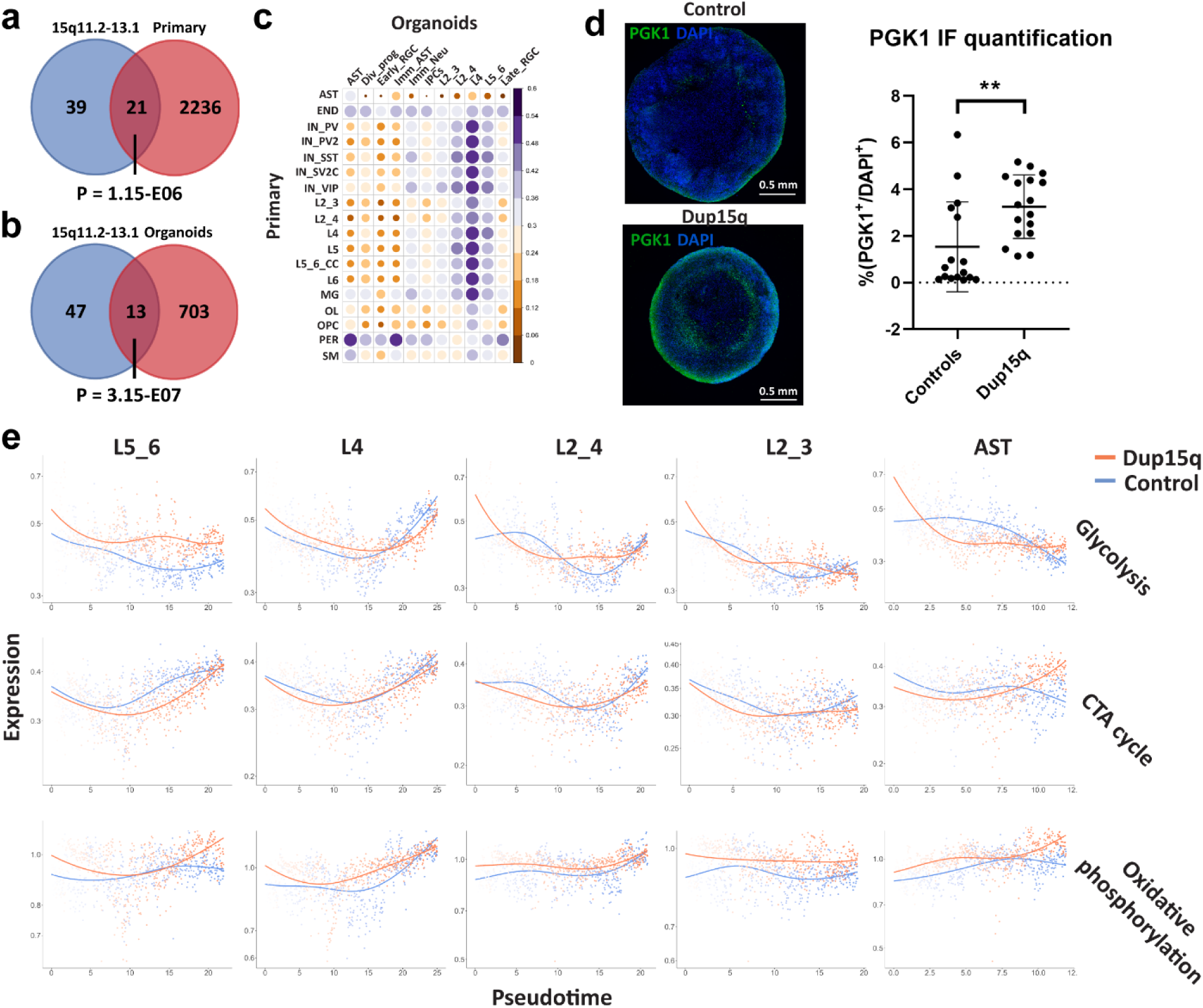
Contribution of duplicated genes to DEGs, cell type gene expression correlation, and metabolic gene set expression of cell-specific trajectories along pseudotime. **a)** Venn diagram showing overlap between putative duplicated genes and primary nuclei overexpressed genes from all cell types (P denotes a hypergeometric p value). **b)** Venn diagram showing overlap between putative duplicated genes and organoid overexpressed genes from all cell types (P denotes a hypergeometric p value). **c)** Pearson’s correlation plot of top 100 expressed genes between organoid and primary cell types. **d)** Immunofluorescent staining of the glycolysis enzyme PGK1 in control and dup15q organoids. The plot on the right is showing quantification summary of PGK1 expression. **e)** Metabolic gene set expression levels of organoid cell-specific trajectories plotted along pseudotime (red line denotes dup15q while blue lines denote control expression).

**Extended Data Figure 6.**
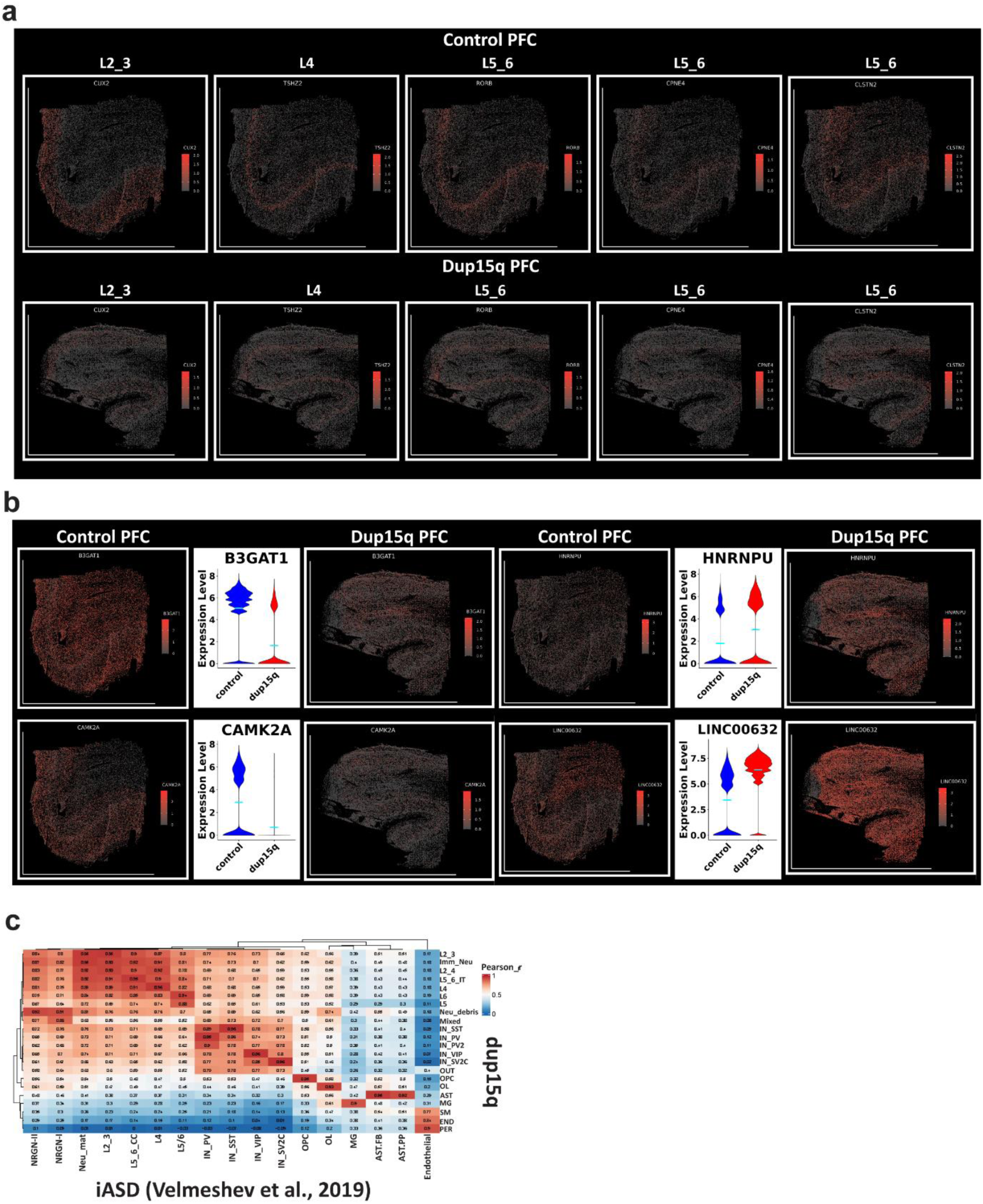
Spatial transcriptomics validates differential gene expression in dup15q syndrome PFC. **a)** Excitatory neuron layer-specific markers tissue expression. **b)** Examples of differentially expressed genes, validated through spatial resolved transcriptomics. **c)** Heatmap with Pearson’s correlation plot of highly expressed genes between dup15q and idiopathic ASD cell types.

## References

1. Sandin, S. et al. The Heritability of Autism Spectrum Disorder. JAMA 318, 1182–1184 (2017).

2. Ben-david, E. & Shifman, S. Networks of Neuronal Genes Affected by Common and Rare Variants in Networks of Neuronal Genes Affected by Common and Rare Variants in Autism Spectrum Disorders. (2014) doi:10.1371/journal.pgen.1002556.

3. Amiri, A. et al. Research article summary. 6720, (2018).

4. Ben-David, E. & Shifman, S. Combined analysis of exome sequencing points toward a major role for transcription regulation during brain development in autism. Mol. Psychiatry 18, 1054–1056 (2013).

5. Parikshak, N. N. et al. XIntegrative functional genomic analyses implicate specific molecular pathways and circuits in autism. Cell 155, 1008 (2013).

6. Voineagu, I. et al. Transcriptomic analysis of autistic brain reveals convergent molecular pathology. Nature 474, 380–386 (2011).

7. Willsey, A. J. et al. XCoexpression networks implicate human midfetal deep cortical projection neurons in the pathogenesis of autism. Cell 155, 997 (2013).

8. Velmeshev, D. et al. Single-cell genomics identifies cell type–specific molecular changes in autism. Science (80-.). 364, 685–689 (2019).

9. Parikshak, N. N. et al. Genome-wide changes in lncRNA, splicing, and regional gene expression patterns in autism. Nature 540, 423–427 (2016).

10. Suzuki, K. et al. Microglial Activation in Young Adults With Autism Spectrum Disorder. JAMA Psychiatry 70, 49–58 (2013).

11. Gandal, M. J. et al. Broad transcriptomic dysregulation occurs across the cerebral cortex in ASD. Nature 611, 532–539 (2022).

12. Li, C. et al. Single-cell brain organoid screening identifies developmental defects in autism. 1–26 (2022).

13. Quadrato, G., et al. Autism genes converge on asynchronous development of shared neuron classes. vol. 602 (Springer US, 2022).

14. Mariani, J. et al. Glutamate Neuron Differentiation in Autism FOXG1-Dependent Dysregulation of GABA / Glutamate. 375–390 (2015) doi:10.1016/j.cell.2015.06.034.

15. Courchesne, E. et al. Neuron Number and Size in Prefrontal Cortex of Children With Autism. 306, 2001–2010 (2011).

16. Goldberg, M. C. et al. Developmental Cognitive Neuroscience Children with high functioning autism show increased prefrontal and temporal cortex activity during error monitoring. Accid. Anal. Prev. 1, 47–56 (2011).

17. Sunkin, S. M. et al. Patches of Disorganization in the Neocortex of Children with Autism. 1209–1219 (2014) doi:10.1056/NEJMoa1307491.

18. Scoles, H. A., Urraca, N., Chadwick, S. W., Reiter, L. T. & Lasalle, J. M. Increased copy number for methylated maternal 15q duplications leads to changes in gene and protein expression in human cortical samples Increased copy number for methylated maternal 15q duplications leads to changes in gene and protein expression in huma. 19, (2011).

19. Matsubara, K. et al. Exploring the unique function of imprinting control centers in the PWS / AS-responsible region : finding from array-based methylation analysis in cases with variously sized microdeletions. 1–7 (2019).

20. Kadoshima, T. et al. Self-organization of axial polarity, inside-out layer pattern, and species-speci fi c progenitor dynamics in human ES cell – derived neocortex. (2013) doi:10.1073/pnas.1315710110.

21. Evolution, H. B. et al. Establishing Cerebral Organoids as Models of Article Establishing Cerebral Organoids as Models of Human-Specific Brain Evolution. Cell 176, 743–756.e17 (2019).

22. Bhaduri, A. et al. Cell stress in cortical organoids impairs molecular subtype specification. Nature 578, 142–148 (2020).

23. Kang, H. J. et al. Spatio-temporal transcriptome of the human brain. Nature 478, 483–489 (2011).

24. Pollen, A. A. et al. Molecular identity of human outer radial glia during cortical development. Cell 163, 55–67 (2015).

25. Uzquiano, A. et al. Proper acquisition of cell class identity in organoids allows definition of fate specification programs of the human cerebral cortex. Cell 185, 3770–3788.e27 (2022).

26. Gandal, M. J. et al. Broad transcriptomic dysregulation occurs across the cerebral cortex in ASD. (2022).

27. Mouse and human share conserved transcriptional programs for interneuron development. 1342, (2021).

28. Nowakowski, T. J. et al. Spatiotemporal gene expression trajectories reveal developmental hierarchies of the human cortex. 1323, 1318–1323 (2017).

29. Iwata, R. & Vanderhaeghen, P. Regulatory roles of mitochondria and metabolism in neurogenesis. Curr. Opin. Neurobiol. 69, 231–240 (2021).

30. Zheng, X. et al. Metabolic reprogramming during neuronal differentiation from aerobic glycolysis to neuronal oxidative phosphorylation. Elife 5, (2016).

31. Ghosh-Choudhary, S., Liu, J. & Finkel, T. Metabolic Regulation of Cell Fate and Function. Trends Cell Biol. 30, 201–212 (2020).

32. Traxler, L. et al. Warburg-like metabolic transformation underlies neuronal degeneration in sporadic Alzheimer’s disease. Cell Metab. 34, 1248–1263.e6 (2022).

33. Mitelman, S. A. et al. Positron emission tomography assessment of cerebral glucose metabolic rates in autism spectrum disorder and schizophrenia. Brain Imaging Behav. 12, 532–546 (2018).

34. Frye, R. E. Mitochondrial Dysfunction in Autism Spectrum Disorder: Unique Abnormalities and Targeted Treatments. Semin. Pediatr. Neurol. 35, 100829 (2020).

35. Vallée, A. & Vallée, J.-N. Warburg effect hypothesis in autism Spectrum disorders. Mol. Brain 11, 1 (2018).

36. Srour, M. et al. Dysfunction of the Cerebral Glucose Transporter SLC45A1 in Individuals with Intellectual Disability and Epilepsy. Am. J. Hum. Genet. 100, 824–830 (2017).

37. Abrahams, B. S. et al. SFARI Gene 2.0: a community-driven knowledgebase for the autism spectrum disorders (ASDs). Mol. Autism 4, 36 (2013).

38. O’Roak, B. J. et al. Multiplex targeted sequencing identifies recurrently mutated genes in autism spectrum disorders. Science 338, 1619–1622 (2012).

39. Gao, Z. et al. An AUTS2–Polycomb complex activates gene expression in the CNS. Nature 516, 349–354 (2014).

40. Pino, E. et al. FOXO3 determines the accumulation of α-synuclein and controls the fate of dopaminergic neurons in the substantia nigra. Hum. Mol. Genet. 23, 1435–1452 (2014).

41. Schäffner, I. et al. FoxO Function Is Essential for Maintenance of Autophagic Flux and Neuronal Morphogenesis in Adult Neurogenesis. Neuron 99, 1188–1203.e6 (2018).

42. Wegiel, J. et al. Significant neuronal soma volume deficit in the limbic system in subjects with 15q11.2-q13 duplications. Acta Neuropathol. Commun. 3, 63 (2015).

43. Fink, J. J. et al. Hyperexcitable Phenotypes in Induced Pluripotent Stem Cell-Derived Neurons From Patients With 15q11-q13 Duplication Syndrome, a Genetic Form of Autism. Biol. Psychiatry 90, 756–765 (2021).

44. Isshiki, M. et al. Enhanced synapse remodelling as a common phenotype in mouse models of autism. Nat. Commun. 5, 4742 (2014).

45. Piochon, C. et al. Cerebellar plasticity and motor learning deficits in a copy-number variation mouse model of autism. Nat. Commun. 5, 5586 (2014).

46. De Rubeis, S. et al. Synaptic, transcriptional and chromatin genes disrupted in autism. Nature 515, 209–215 (2014).

47. Satterstrom, F. K. et al. Large-Scale Exome Sequencing Study Implicates Both Developmental and Functional Changes in the Neurobiology of Autism. Cell 180, 568–584.e23 (2020).

48. Velmeshev, D. et al. Single-cell genomics identifies cell type-specific molecular changes in autism. Science 364, 685–689 (2019).

49. Stuart, T. et al. Comprehensive Integration of Single-Cell Data. Cell 177, 1888–1902.e21 (2019).

50. Butler, A., Hoffman, P., Smibert, P., Papalexi, E. & Satija, R. Integrating single-cell transcriptomic data across different conditions, technologies, and species. Nat. Biotechnol. 36, 411–420 (2018).

51. Caglayan, E., et al. Ambient RNA analysis reveals misinterpreted and masked cell types in brain single-nuclei datasets. (2022).

52. Finak, G. et al. MAST: a flexible statistical framework for assessing transcriptional changes and characterizing heterogeneity in single-cell RNA sequencing data. Genome Biol. 16, 278 (2015).

53. Cao, J. et al. The single-cell transcriptional landscape of mammalian organogenesis. Nature 566, 496–502 (2019).

54. Liberzon, A. et al. The Molecular Signatures Database Hallmark Gene Set Collection. Cell Syst. 1, 417–425 (2015).

55. Morabito, S., Reese, F., Rahimzadeh, N., Miyoshi, E. & Swarup, V. High dimensional co-expression networks enable discovery of transcriptomic drivers in complex biological systems. 1–32 (2022).

56. Kuleshov, M. V et al. Enrichr : a comprehensive gene set enrichment analysis web server 2016 update. 44, 90–97 (2016).

57. Mi, H., Muruganujan, A., Casagrande, J. T. & Thomas, P. D. Large-scale gene function analysis with the PANTHER classification system. Nat. Protoc. 8, 1551–1566 (2013).

## Methods References

1. Sandin, S. et al. The Heritability of Autism Spectrum Disorder. JAMA 318, 1182–1184 (2017).

3. Amiri, A. et al. Research article summary. 6720, (2018).

23. Kang, H. J. et al. Spatio-temporal transcriptome of the human brain. Nature 478, 483– 489 (2011).

